# Divergent Dopamine Dynamics within the Accumbens Core

**DOI:** 10.64898/2026.09.23.753606

**Authors:** Seongtak Kang, Dennis A. Burke, Caroline Kornbrek, Farah Farouq, Huijeong Jeong, Vijay Mohan K. Namboodiri, Joshua D. Berke

## Abstract

Dopamine in the nucleus accumbens Core powerfully influences motivation and learning, yet the information encoded by Core dopamine remains unresolved. Competing theories typically assume a spatially uniform signal, and standard recording methods average dopamine over substantial tissue volumes. Using two-photon imaging through microprisms in behaving mice, we reveal a pronounced spatial organization of Core dopamine dynamics. Posterior Core showed phasic dopamine increases to both rewards and aversive stimuli, with transfer to a predictive cue after learning. Anterior Core showed phasic decreases to aversive stimuli, weaker increases to reward, and higher tonic dopamine that was further amplified by cocaine. Rather than uniform or highly fragmented signals, the Core contains a smoothly graded mixture between two distinct sets of dopamine dynamics, neither matching canonical reward prediction errors.

## Main Text

The nucleus accumbens (NAc) is a key neural hub for motivated behavior (*1*). Dopamine in the NAc - especially its “Core” region - is critical for the willingness to work to obtain rewards (*2*, *3*) and for learning associations between events and rewards (*4*). Addictive drugs enhance NAc dopamine release, and dysregulated NAc dopamine is implicated in multiple psychiatric conditions including mood disorders (*5*).

Despite decades of investigation, the specific information conveyed by NAc dopamine, and the spatial and temporal scales of these signals, remain unclear. NAc receives input from ventral tegmental area (VTA) dopamine cells, whose firing often resembles a reward prediction error (RPE; (*6*)). RPE-like activity includes brief (“phasic”) firing increases after unexpected rewards, transfer of reward responses to reward-predictive cues after learning, and firing decreases after reward omission or punishments (*7–9*). However, individual VTA dopamine cells also show considerable heterogeneity in their functional activity patterns (*10*, *11*) and projection targets (*12*). The extent to which such heterogeneous activity results in spatiotemporally complex patterns of dopamine release is unknown.

Dopamine has been shown to have distinct dynamics, and functional impact, in NAc Core vs. the adjoining Shell (*13–16*). But it has been broadly assumed that dopamine signals are spatially uniform within the Core itself. This assumption reflects limitations of spatial resolution: e.g. the widely used fiber photometry approach samples fluorescence from a poorly defined volume hundreds of microns wide. Similarly, limitations of temporal resolution and/or stability of dopamine recordings have constrained our understanding of slower (“tonic”) dopamine changes (*8*, *17*).

Perhaps because of these constraints, existing measurements of Core dopamine have not yielded a consistent picture. Dopamine release in the Core often shows phasic increases that scale with RPE, and phasic decreases following aversive events (*14*, *18*). However other studies report that Core dopamine increases with aversive events (*19*, *20*). Core dopamine increases are also often coupled to movement initiation and reward approach (*17*, *21–23*), and can persist for many seconds, longer than expected for an error signal (*21*). Such observations have led to accounts in which Core dopamine conveys alternative quantities such as value, perceived salience, causal meaningfulness, or learning rate (*17*, *20*, *24–26*). Even more prolonged changes in NAc dopamine (*27*) have been hypothesized to be a separate, motivation-related signal (*28–30*), for example encoding reward rate (*31*).

The actual granularity of dopamine – i.e. the number of distinct dopamine signals – has strong computational implications. Most learning-related dopamine theories posit widely broadcast scalar signals. But in some recent formulations, dopamine instead provides detailed (“vectorized”) error signals that may enable the brain to learn not just predictions of aggregate future reward, but also the specific identity and timing of upcoming events (*32–38*). This would require variation between VTA dopamine cells to be preserved as complex local patterns of downstream dopamine release, rather than blended through diffusion into a scalar RPE (*39*). To arbitrate between current theories, we need to determine the dimensionality of dopamine: is there just one spatially homogeneous Core dopamine signal, a mixture of a small set of distinct signals, or a rich pattern of many signals?

Here we use chronic deep microprism implants (*40*), together with the advanced fluorescent dopamine sensor dLight3.8 (*41*), to simultaneously image dopamine dynamics over a wide span of the Core with high spatial and temporal resolution. We find that anterior and posterior Core diverge sharply in their tonic dopamine levels, spontaneous fluctuations, and phasic responses to rewarding and aversive stimuli. Contrary to existing theories, Core dopamine appears organized as neither a single homogeneous field, nor a collection of many distinct signals, but as a graded mixture of approximately two distinct sets of dynamics.

### Two-photon imaging of Core dopamine

To visualize dopamine dynamics, we infused adeno-associated virus for broad expression of dLight3.8 into the NAc of adult mice and chronically implanted a microprism lateral to NAc (**Fig. 1A**). The microprism face provided optical access to a ≈1.5 × 1.5 mm sagittal field of view (FOV) spanning the anterior-posterior extent of the Core (note that anterior Core is also more dorsal to posterior Core). Each implant remained stable for months, enabling longitudinal imaging of the same FOV across behavioral conditions and pharmacological manipulations.

**Figure 1:**
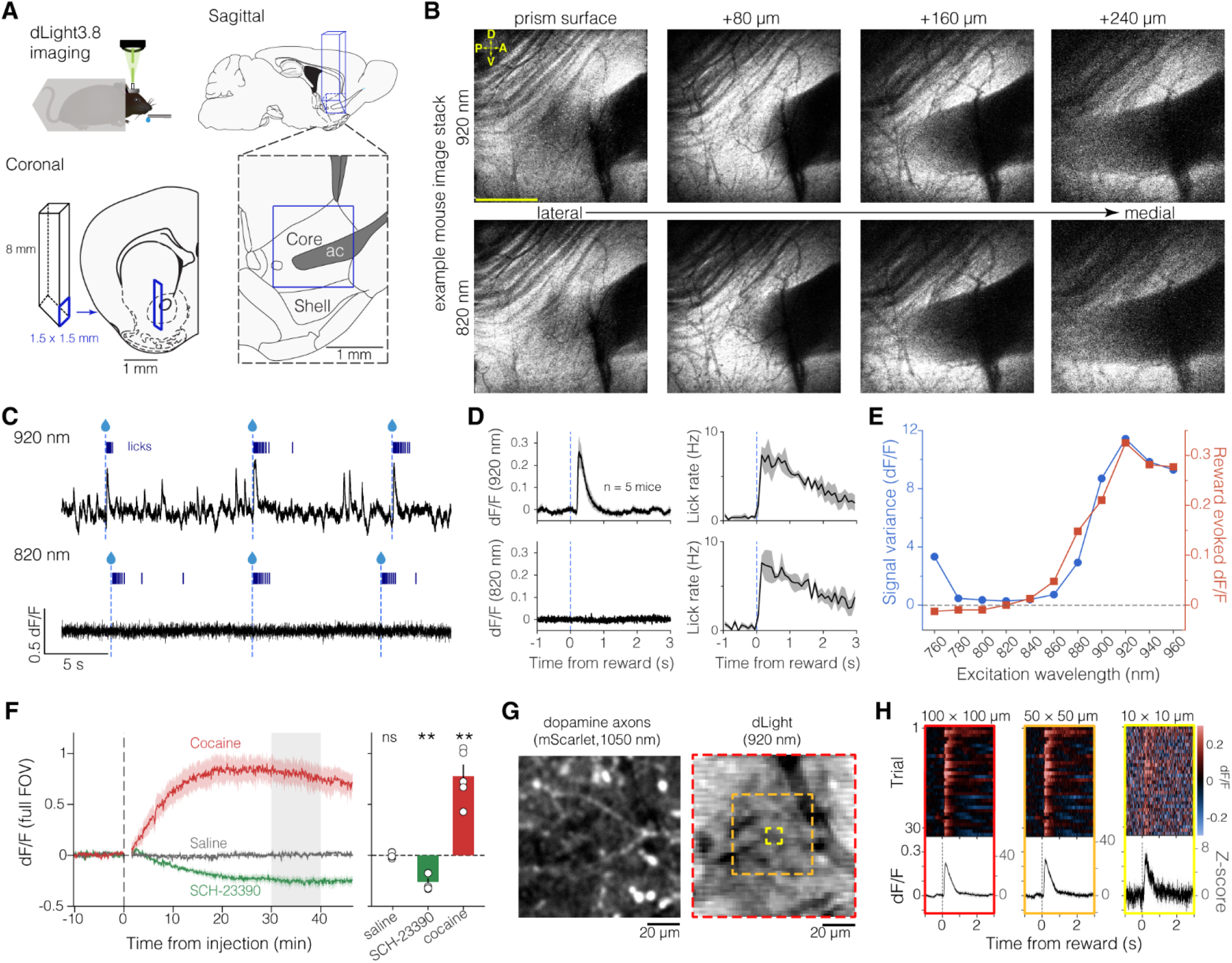
High-resolution imaging of NAc Core dopamine. **A.** Schematic of two-photon imaging of accumbens dopamine in awake, behaving mice. Microprism was implanted with the imaging window facing medially providing a sagittal view of Core. Atlas section (Franklin & Paxinos, 2008) shows prism field-of-view (FOV) corresponding to the example mouse in **B.** ac: anterior commissure. **B.** Imaging “Z”-stack for example mouse (Mouse 3). Each image shows the full FOV (≈1.42 mm by ≈1.42 mm) at multiple focal depths, using either 920 nm (top) or 820 nm (bottom) excitation and averaging 512 frames (acquired at ≈30 Hz with 512 × 512 pixels; ≈2.8 µm/pixel). Due to the prism mirror, focusing deeper into the tissue corresponds to imaging more medially. Distances noted on each column refer to physical distances traveled by microscope, without adjusting for the refractive index of the glass prism. In each image, contrast was auto-scaled from the 5th to the 95th percentile of pixel intensities. Scale: 500 µm. Orientation: D, dorsal; V, ventral; A, anterior; P, posterior). **C-E**. Dependence of dLight3.8 signal on excitation wavelength. (**C**) Average full-FOV dF/F traces from example mouse (Mouse 1) recorded at 920 nm (top) and 820 nm (bottom) excitation (64 × 64 pixels, ≈22 µm/pixel, ≈205 Hz) during delivery and consumption of unpredicted sucrose solution (15% w/v) rewards (mean inter-reward interval = 15 s). Vertical dashed blue lines denote reward delivery. Blue ticks above traces denote licks at reward delivery spout. (**D**) Mean (n = 5 mice) fluorescence (left) and licking behavior (right) aligned to reward delivery (blue dashed line), using either 920 nm (top) or 820 nm (bottom) excitation (20 rewards per wavelength). Error shading represents SEM. (**E**) Reward response (mean of signal 0 – 0.5 s following reward delivery) and spontaneous signal variance were greatest at 920 nm and minimized at 820 nm (n = 2 mice, 20 rewards / wavelength). **F.** Effects of systemic drugs on dLight signal (920nm excitation), comparing saline (10 ml/kg, gray), SCH-23390 (0.3 mg/kg), and cocaine (red; 10 mg/kg). Mice were awake and received sporadic rewards (mean inter-reward interval = 75 s). *Left,* mean time course of signal change from baseline (n = 5 mice, shading = SEM). Data were not recorded during drug injections. *Right,* Average signal change from baseline 30-40 minutes after injection (gray shading on left). Circles represent individual mice; error bars, SEM. Saline: dF/F = 0.007 ± 0.012, one-sample t-test: t(4) = 0.55, p = 0.61; SCH-23390: dF/F = −0.295 ± 0.037, t(4) = −7.99, p = 0.0013; cocaine: dF/F = 0.697 ± 0.109, t(4) = 6.37, p = 0.0031. ns: not significant. Note that all statistical tests were two-tailed. **G.** Dopamine axons expressing mScarlet (left, 1050 nm excitation) and dLight (right, 920nm) within a local zone inside the Core, imaged in Mouse 4. Subsequent analysis used the regions-of-interest (ROIs) demarcated in red (100 x 100 µm), orange (50 x 50 µm) and yellow (10 x 10 µm). **H**. dLight signal changes around reward (n = 32 rewards, mean IRI = 15 s) for the ROIs shown in G. *Top,* individual rewards; *bottom,* averages. Scales indicate dF/F amplitude (left axis) and Z-score (right axis). A smaller ROI leads to a more variable signal (100µm: baseline SD = 0.0067, 50µm: baseline SD = 0.0093, 10µm: baseline SD = 0.0403), but the reward response remains consistently detectable.

The Core encircles the anterior commissure (*42*) and is bounded ventrally and medially by the Shell, which is neurochemically distinct (e.g. calbindin-poor; (*43*)). Dorsally, the Core merges into the rest of striatum without a specific boundary (*44*, *45*). Adjustable focal depth allowed imaging over a ∼300µm medial-lateral range (**Fig. 1B; fig. S1**); in each mouse, we chose an imaging plane to maximize dLight fluorescence, within 150µm of the anterior commissure. Histology including calbindin labelling confirmed that prism FOVs included both anterior and posterior portions of the Core in all mice (**fig. S2**).

The sensitivity of dLight3.8 to dopamine depends on excitation wavelength (*41*). To test this sensitivity under our two-photon, *in vivo* conditions, we serially imaged at different wavelengths while presenting mice with unpredictably timed rewards (15% sucrose water drops). With 920 nm excitation, fluorescence averaged across the FOV was highly dynamic, with robust increases following reward delivery (**Fig. 1C, D**). With 820 nm excitation, no spontaneous or reward-related changes were seen, despite mice avidly licking after rewards (**Fig. 1C, D**). To systematically characterize this wavelength dependence, in two mice we swept the excitation from 760 to 960 nm in 20nm steps. Both reward-evoked increases, and the signal variance associated with spontaneous dopamine fluctuations, peaked at 920 nm and fell to near zero around 820 nm (**Fig. 1E, fig. S3**). Thus, under these recording conditions, 920 nm excitation provides the most dopamine-sensitive signal, while 820 nm serves as an effective isosbestic (dopamine-independent) signal. This contrast allows us to distinguish spatial variation in dopamine from spatial variation in dLight expression.

We further confirmed the dopamine-dependence of 920 nm excitation signals using pharmacological manipulations. The dopamine D1 receptor antagonist SCH-23390 (0.3 mg/kg, s.c.) produced a clear, sustained decrease in FOV-averaged fluorescence (**Fig. 1F**), consistent with the drug competing with endogenous dopamine for dLight binding. Conversely, the dopamine reuptake inhibitor cocaine (10 mg/kg, s.c.), known to increase Core dopamine (*46*, *47*), produced a large, sustained increase in mean fluorescence (**Fig. 1F**).

To confirm the capability of our imaging setup to resolve fine spatial structure, in two mice we also expressed a red fluorophore in dopamine axons (by infusing virus for Cre-dependent expression of mScarlet into the VTA of DAT-Cre mice). This revealed individual dopamine axons within Core (**Fig. 1G**). We then examined whether the dLight signal is sufficiently robust to detect spatially fine dopamine fluctuations, by comparing regions of interest (ROIs) with widths ranging from 100µm down to 10µm (**Fig. 1G,H**). As expected, signal:noise was higher when averaging more pixels, but even with 10µm ROIs, we could readily detect the dopamine response to individual rewards, and the mean dF/F response was similar across spatial scales (**Fig. 1H**). Having thus established that dLight imaging can report dopamine release with high spatial resolution, we could critically assess the spatial homogeneity of Core signals.

### Greater tonic dopamine in anterior Core

We first assessed whether average dopamine levels are spatially uniform within Core. For each mouse, we compared dopamine-sensitive 920nm images to dopamine-insensitive 820nm images for the same FOV (**Fig. 2A)**, using “baseline” recordings without any experimental manipulation. This revealed higher mean dopamine signals in the anterior/dorsal portion of the FOV compared to the posterior/ventral portion, for all five mice (**Fig. 2B,C, fig. S4**). This anatomical difference was apparent even when comparing pixels with similar 820nm fluorescence – i.e. controlling for dLight expression (**fig. S5)**. For further quantification, in each mouse we selected one “anterior” and one “posterior” ROI (≈265 × 265µm), both within the Core, choosing, based on histology, areas that best matched locations across animals (**Fig. 2B, fig. S1**). The anterior Core ROI showed consistently higher average dopamine (**Fig. 2C**, right). In a subset of the same mice (n = 4), we also imaged dLight using two-photon Fluorescence Lifetime Imaging (FLIM) (*41*, *48*). This complementary approach, also known to be relatively insensitive to fluorophore expression or non-specific drift (*49*, *50*), confirmed greater average dopamine levels in anterior, compared to posterior Core (**fig. S6**).

**Figure 2:**
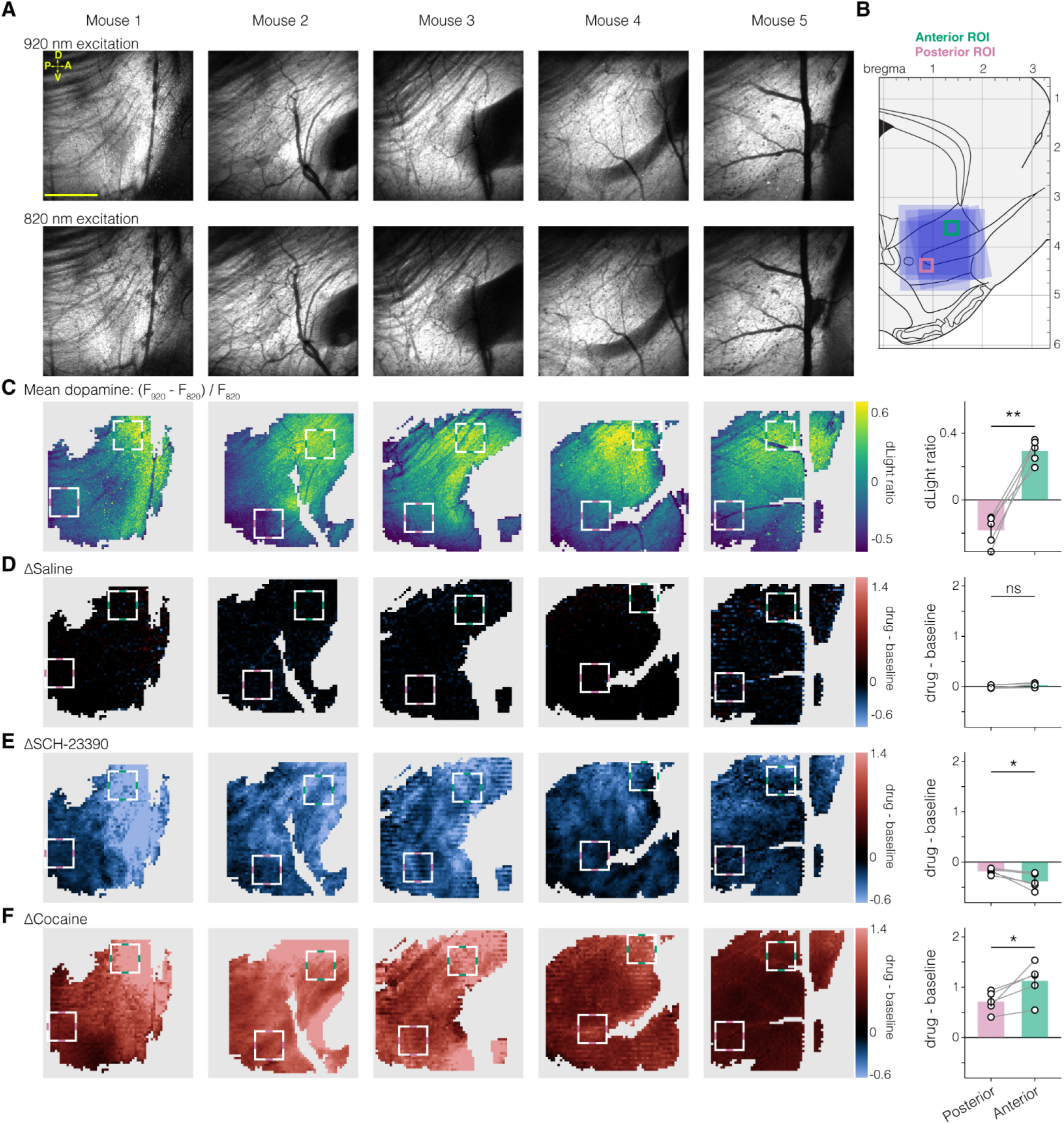
Elevated dopamine levels in anterior Core. **A.** Dopamine-dependent (top, 920 nm excitation) and dopamine-independent (bottom, 820nm) mean images for each mouse (100s recording, ≈30 Hz, 512 x 512 pixels). Images at each wavelength were recorded sequentially in the same session. Scale bar: 500 µm. **B.** Positions of each prism FOV relative to sagittal brain atlas section (Franklin & Paxinos, 2008) (see also **figs. S1, S2**). Each blue square represents one mouse’s FOV. Pink and green squares represent anatomically consistent (in the anterior-posterior and dorsal-ventral axes) 265 × 265 µm anterior (green) and posterior (pink) ROIs used in this and subsequent figures. Axis numbers indicate distance from bregma in mm. **C.** Mean dopamine maps. *Left,* Normalizing the 920 nm image by the dopamine independent 820 nm image (“dLight ratio”: (920 nm – fitted 820 nm) / fitted 820 nm) shows a stronger signal in anterior regions. FOVs were masked before analysis to remove pixels with low 820 nm signal (**fig. S4**, see Methods). *Right,* Mean dLight ratio in anterior (green) and posterior (pink) ROIs (anterior +0.293 ± 0.031, posterior −0.183 ± 0.040; paired t-test, n = 5 mice, t(4) = 8.10, p = 0.0013). **D-F.** *Left,* Difference maps showing drug-induced changes in dLight ratio (same experiment as Fig. 1F). All maps share the same color scale (at right). *Right,* Quantification and comparison for anterior (green) and posterior (pink) ROIs. Paired t-tests (n = 5 mice): saline: t(4) = 1.91, p = 0.13 (n.s.); SCH-23390, t(4) = −3.15, p = 0.035; cocaine: t(4) = 3.17, p = 0.034).

In zones with higher dopamine, blocking the dLight dopamine-binding site should produce a greater decrease in 920nm-excited fluorescence. Indeed, anterior Core showed a more pronounced decrease in dLight signal following administration of the competitive dopamine antagonist SCH-23390, compared to posterior Core (**Fig. 2D,E**; see **fig. S7** for additional details on each mouse). Together, these findings indicate that anterior Core has higher average dopamine release. This higher level did not saturate the dLight3.8 sensor, as anterior Core showed especially strong dopamine increases after cocaine (**Fig. 2D,F, fig. S7**), potentially indicating greater sensitivity of this subregion to drugs of abuse.

A higher time-averaged dopamine level could arise from greater “tonic” (continuous, or slowly varying) levels, or a greater frequency of “phasic” (transient) release events (*2*, *31*). Although phasic events have typically been assumed uniform within the Core, we found that spontaneous activity was clearly non-uniform (**Fig. 3A,B**, see **fig. S8** for additional examples in each mouse). While some fast dopamine changes involved most of the FOV, many increases and decreases were more spatially selective **(Video 1)**. As an initial characterization, we assessed whether there is a consistent anatomical axis along which this spatiotemporal variation is most prominent. We used a spectral clustering algorithm (*51*, *52*) to force each mouse’s FOV into two clusters based solely on spontaneous dopamine dynamics **(Fig. 3C)**. In every mouse, the spatial locations of these two clusters were anterior/dorsal vs posterior/ventral, encompassing our previously defined anterior and posterior Core ROIs. This demonstrates a consistent axis of variation for Core phasic dopamine events (but note it does not indicate that there are exactly two dopamine zones separated by a clear boundary).

**Figure 3:**
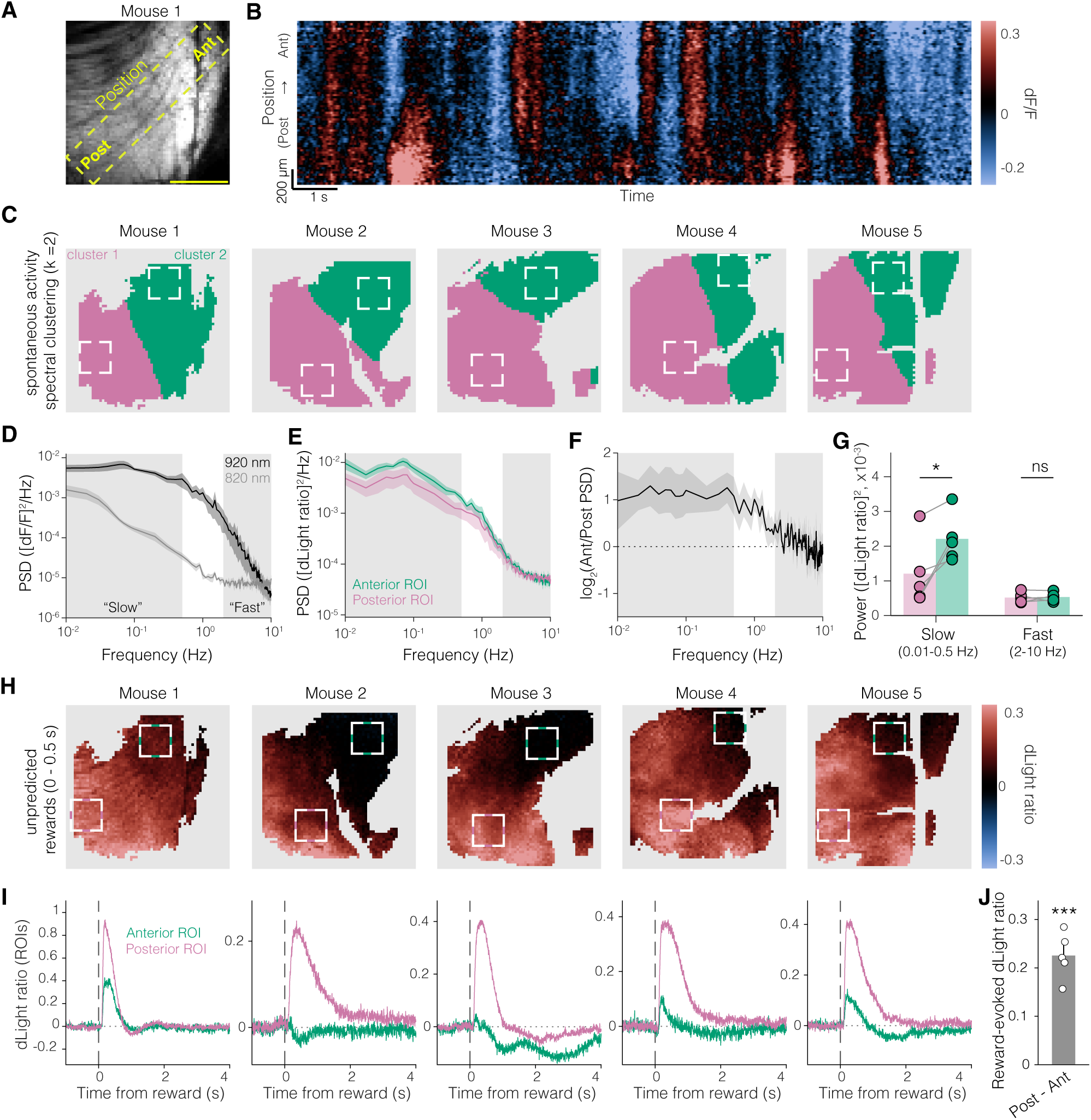
Distinct phasic dopamine dynamics in anterior and posterior Core. **A.** FOV for example Mouse 1 overlaid with 15-pixel wide diagonal ROI from posterior (“Post”) to anterior (“Ant”) Core. Scale bar, 500 µm. **B.** Kymograph of example spontaneous changes in dLight signal along the anterior-posterior diagonal marked in **A**. Throughout this figure, images were acquired at ≈205 Hz, 64 × 64 pixels (≈22 µm/pixel). **C.** Unbiased spectral clustering of spontaneous activity (20 min; see *Methods*) into two clusters (k = 2) consistently separates posterior/ventral (pink) and anterior/dorsal (green) portions of the Core. Squares represent the same anterior and posterior ROIs as before (Fig. 2B). **D.** Full-FOV absolute power spectra of the 920 nm (dopamine-dependent) and 820 nm (dopamine-independent) signals during sessions with unpredicted reward deliveries (n = 5 mice, 32 rewards each, 15 s mean inter-reward interval). Spectra converge at high frequencies, but dopamine-dependent signals are higher over a wide frequency range. Gray shading marks “slow” (0.01–0.5 Hz) and “fast” (2–10 Hz) bands. **E.** Absolute power spectra for spontaneous activity in anterior (green) and posterior (pink) ROIs. Shading, SEM across mice. **F.** Log₂ ratio of anterior to posterior ROI power spectra (> 0 implies greater anterior power). Shading, SEM. **G.** Absolute band power, anterior vs posterior. Paired t-test within each mouse (n=5), per band: slow, t(4) = 3.6, p = 0.023; fast, t(4) = 0.289, p = 0.787 (n.s.). Error bars, SEM. **H.** Maps of unpredicted reward response in each mouse (mean dLight ratio 0–0.5 s after reward delivery). Maps were averaged over two recording sessions (64 rewards each; mean inter-reward intervals of 15 and 75 s). Squares, anterior/posterior ROIs as before. **I.** Mean reward response time course in anterior (green) and posterior (pink) ROIs. **J.** Comparing reward response by posterior-minus-anterior dLight ratio (0–0.5 s post-reward, n=5 mice). One-sample t-test vs zero, t(4) = 10.49, p=0.0005.

Since anterior and posterior Core can have distinct phasic dopamine release, we examined whether a greater rate of phasic fluctuations in anterior Core may be responsible for the higher time-averaged dopamine noted above. Instead, anterior Core showed similar power at higher frequencies associated with phasic dopamine, but higher power at low frequencies (**Fig. 3D-G**). Taking these results together, we conclude that the elevated time-averaged baseline signal in anterior Core reflects greater tonic dopamine levels, rather than more frequent phasic events.

### Reward-to-cue response transfer occurs preferentially in Posterior Core

We next turned from spontaneous to event-locked dopamine signals. Unpredicted rewards increased FOV-average dopamine signals, as noted earlier, but surprisingly this response was not spatially uniform (**Fig. 3H).** Across all mice, reward responses in posterior Core were stronger, while those in the anterior Core were weaker with higher variability between animals **(Fig. 3I,J; Video 2)**.

A seminal observation is the temporal transfer of dopamine reward responses to earlier predictors, consistent with dopamine signaling prediction errors. We therefore assessed how evoked dopamine patterns changed as mice learned a Pavlovian cue–reward task. Two distinct tones (CS+ or CS-, each 250 ms duration) were followed 2 s later by reward or nothing respectively. On the first day of conditioning, animals showed no anticipatory licking (**Fig. 4A**) or dopamine responses to cues; dopamine reward responses were larger in Posterior Core as noted before **(Fig. 4B)**. Over 2 weeks of training, mice developed robust anticipatory licking to the CS+ **(Fig. 4C, E**). As expected, this conditioned behavior was associated with a clear CS+ dopamine response when averaging over the full FOV **(Fig. 4C, E**). However, this CS+ response was driven preferentially by posterior Core **(Fig. 4C-F; Video 3**); CS+ responses in anterior Core were weaker, less consistent and on average not significantly different from zero (**Fig. 4C-F**; see **fig. S9** for dopamine maps in each mouse). After learning, reward responses were comparable in posterior and anterior Core (**Fig. 4E, F**). The CS-cues evoked minimal anticipatory licking **(Fig. 4G)**, only a small posterior dopamine response, and no significant anterior response **(fig. S10)**.

**Figure 4:**
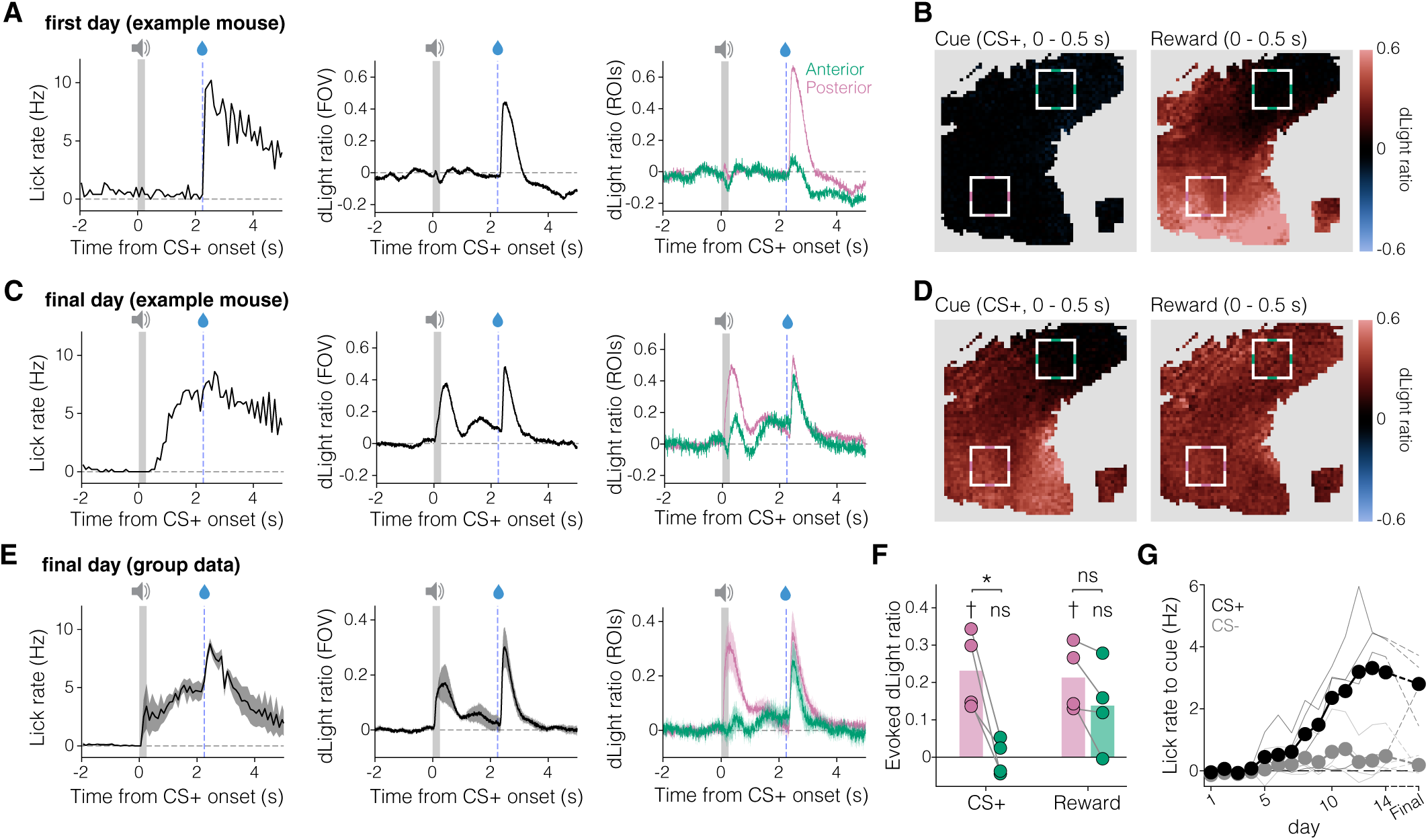
Reward-to-cue transfer specifically in posterior Core. **A.** Example mouse (Mouse 3) on the first day of cue-reward conditioning. *Left,* mean CS+ aligned licking (*left*), dLight over the full FOV (*middle*), and dLight in anterior (green) and posterior (pink) ROIs (*right*). Included are all 50 CS+ trials per day. Gray shading represents CS+ duration. Dashed blue line denotes reward delivery. For CS-results, see **fig. S10**. **B.** Corresponding first-day dopamine maps for the same example mouse, averaging dLight data after CS+ onset (0–0.5 s; *left*) or reward delivery (0–0.5 s; *right*). **C.** Final day of cue-reward conditioning for the same example mouse with data presented as in **A**. **D.** Corresponding final day dopamine maps for the same example mouse. **E.** Final day group data (n = 4 mice; shading, SEM). **F.** Quantification of dopamine responses to CS+ (0–0.5 s) and reward (0–0.5 s) in anterior (green) and posterior (pink) ROIs. One-sample t-tests (n = 4 mice) comparing to zero: posterior CS+, t(3) = 4.396, p = 0.022 (†); anterior CS+, t(3) = −0.04, p = 0.971 (n.s.); posterior reward, t(3) = 4.70, p = 0.018 (†); anterior reward, t(3) = 2.37, p = 0.098 (n.s.). Anterior and posterior ROIs were also compared to each other: paired t-tests: CS+, t(3) = 4.465, p = 0.021 (*); reward, t(3) = 2.376 p = 0.098 (n.s.). **G.** Licking to CS+ and CS-over learning. Lick rates are measured in the 2.25 s following cue onset, and baseline subtracted. Circles and thicker lines represent mean across mice; thin lines represent individual mice.

### Opposite responses to aversive stimuli in Anterior vs Posterior Core

A strong dopamine response to unpredicted rewards, together with transfer to a predictive cue, appears consistent with RPE-like coding in posterior, but not anterior, Core. To assess this further, we used aversive stimuli. In canonical RPE theories, aversive events are either irrelevant or counted as negative rewards, and should therefore produce absent or negative dopamine responses. According to other theories, dopamine should increase to a broad range of unexpected salient or meaningful events, including aversive stimuli (*20*, *26*, *53–57*). Prior studies recording dopamine release without fine spatial resolution are inconsistent: both positive and negative Core responses have been reported (*14*, *18–20*, *58–60*).

Using an infrared laser, we delivered brief (200 ms) noxious heat pulses to the whisker pad (**Fig. 5A**) in four mice. Each mouse displayed reliable, time-locked unconditioned behavioral responses (“face wiping”, **Fig. 5B**) (*61*). In contrast to this behavioral consistency, the dopamine response averaged over the full FOV varied markedly: two mice showed net phasic increases, and two showed net phasic decreases (**Fig. 5B**). This inter-animal heterogeneity mirrors the variability across past studies.

**Figure 5:**
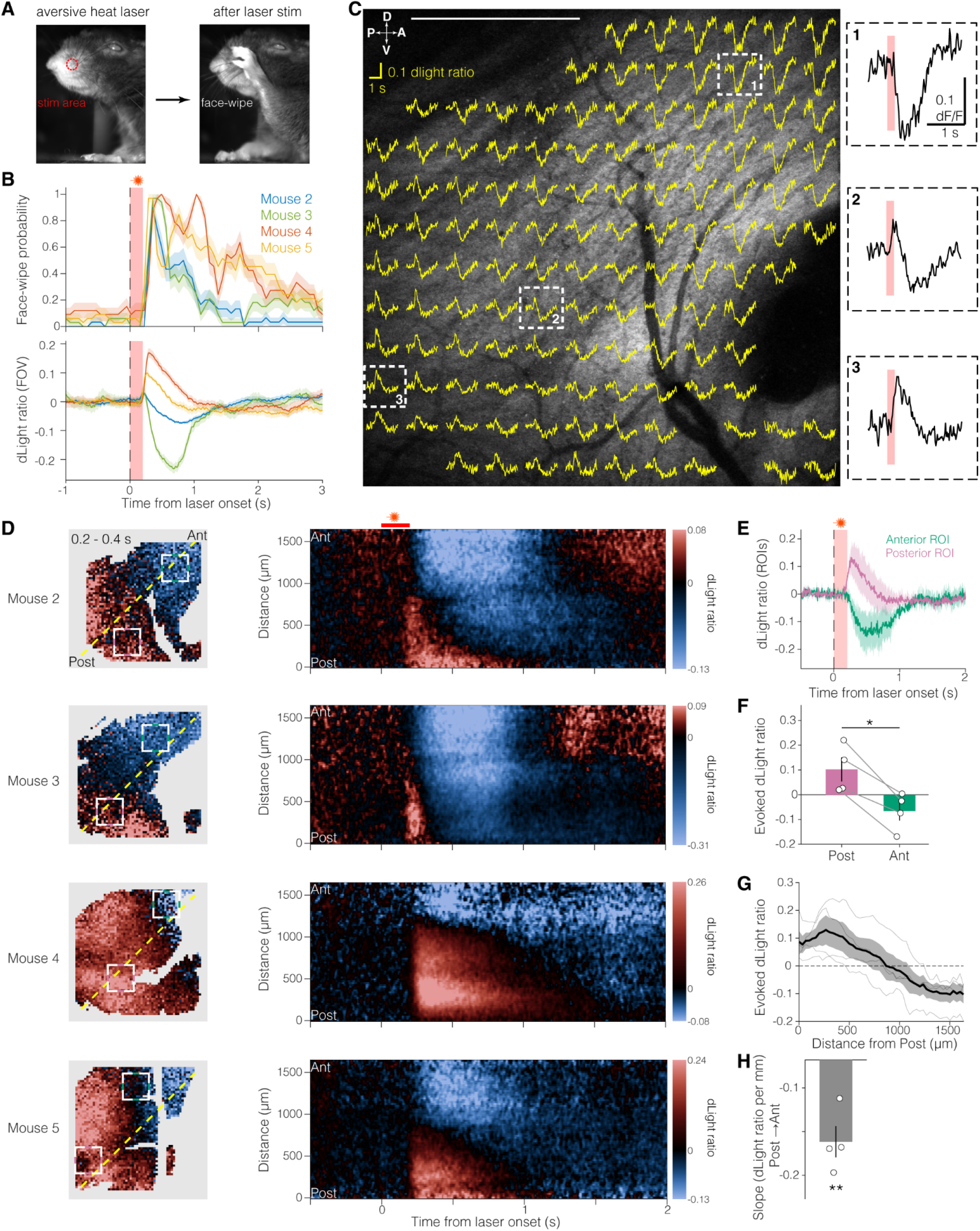
Opposing aversive dopamine responses within Core. **A.** *Left,* infrared laser (1450nm) was directed at each mouse’s whisker pad (200 ms pulses; 30 pulses per session, 60 s mean inter-stimulus interval). *Right,* face-wipe response. **B.** Heat stimulus increases face-wipe probability in all mice tested (*top*; vertical paw movement, see Methods), but the full-FOV Core dopamine response is variable (*bottom*) aligned to laser onset. Shading, SEM across trials for each mouse. **C.** Map of dopamine time courses in response to heat stimulus, in example mouse (Mouse 2). The full FOV was divided into a grid of 5 × 5 pixel zones (≈22 µm/pixel, recorded at ≈205Hz). Yellow traces show mean dLight signals for each zone (−0.5 – 2s relative to heat onset). White dashed boxes indicate three illustrative zones, enlarged on the right (red, laser stimulation time). For anatomical illustration, traces are superimposed on a higher-resolution (512 x 512) image for the same FOV. **D.** *Left,* Maps of early (0.2 – 0.4s) dopamine response to heat stimulus in each mouse. Squares, same anterior and posterior ROIs as before. Dashed line represents the center of the diagonal band (15 pixels wide) used for kymographs. *Right,* kymographs of dopamine response in each mouse. **E.** Mean time course of dopamine response in anterior (green) and posterior (pink) ROIs. Shading, SEM across mice (n = 4). **F.** Mean dopamine response (0.2 – 0.4 s after heat onset) in anterior (green) and posterior (pink) ROIs. Error bars, SEM across mice (n = 4). Paired t-test, anterior vs. Posterior: t(3) = 5.273, p = 0.013. **G.** Gradient of mean dopamine response (0.2 – 0.4 s after heat onset) by distance along the FOV diagonal (dashed line in D), from posterior/ventral (Post) to anterior/dorsal. Shading, SEM across mice; thin lines, individual mice. **H.** Slopes of the dopamine response gradients (0.2 – 0.4 s after heat onset) shown in G (linear least-squares fit). Slopes were significantly negative (one-sample t-test against zero: t(3) = −9.118, p = 0.0028).

Spatially resolved imaging cleared up this apparent inconsistency. Within each animal, aversive stimulation evoked *both* dopamine increases and decreases, in posterior and anterior regions of the FOV respectively (**Fig. 5C-F; fig. S11; Video 4)**. Inspection of the dopamine time course within local zones (**Fig. 5C, fig. S11)** showed a smoothly graded transition from very fast increases (posterior) to fast decreases (anterior); in between, the dopamine response resembled a mixture of both responses. Quantification across all four mice, using the same ROIs as before, confirmed that this aversive stimulus evoked a positive dopamine response in posterior Core and a negative response in anterior Core (**Fig. 5E-H**). Neither behavior nor dopamine response consistently declined across trials, arguing against purely novelty-related saliency signals (*20*) (**fig. S12**).

Overall, in both rewarding and aversive contexts dopamine dynamics show dramatic differences between anterior and posterior Core, with neither zone showing patterns expected from conventional RPE theory.

## Discussion

By combining microprism-based two-photon imaging, the enhanced dopamine sensor dLight3.8, and a series of independent behavioral and pharmacological tests, we have shown that dopamine dynamics within the accumbens Core are organized along an anterior/dorsal - posterior/ventral axis. Anterior Core dopamine is characterized by greater tonic levels, stronger enhancement by acute cocaine, strong phasic decreases to aversive stimuli, and weak phasic increases to rewards. Posterior Core, by contrast, displays robust phasic increases to both rewards and aversive stimuli, with transfer of responses to reward-predictive cues.

How many dopamine signals are there? Classic theory posited one: a global RPE precise in time, but uniform over all forebrain targets (*9*). Subsequent measurements of bulk dopamine release showed differences between targets, including striatal subregions separated on a scale of mm (*13–16*, *62–66*). Here we have revealed a finer spatial scale to NAc dopamine: distinct dynamics even within the Core. It is possible that yet finer scales exist (*67–69*), so that the number of distinct Core dopamine signals is very large. However, our results constrain this possibility. Between adjacent locations in Core, mean dopamine responses varied smoothly, even when using small (≈20µm wide) ROIs. This is consistent with prior imaging studies in dorsal striatum: even though adjacent individual dopamine axons can differ strongly in activity (*70*), net dopamine release is highly correlated between sites separated by hundreds of µm (*71*). This also fits the broad axonal fields of individual dopamine cells, each spanning hundreds of µm and overlapping such that a given striatal neuron may be innervated by ∼100 distinct dopamine cells (*72*). Overall, Core dopamine seems best described as a smooth spatial gradient between two characteristic sets of dynamics, rather than a single uniform signal, or two separate zones with a distinct boundary, or a highly fragmented mosaic of microdomains. Nonetheless, further work is certainly needed to explore both very fine spatial scales (such as individual dopamine release sites) and the broader distribution of dopamine dynamics throughout NAc and beyond in behaving animals. There may also be important features of dopamine signals that are not visible in our experiments using dLight3.8 - e.g. because the sensor affinity constrains the detectable range of dopamine concentrations.

What do the two divergent dopamine dynamics encode? Response transfer to cues from rewards, as observed in posterior Core, is a key feature of many learning-related dopamine models (*24*, *26*, *73*); though not all (*20*). However, the increased dopamine to aversive stimuli in posterior Core is inconsistent with canonical RPE coding. It is more consistent with theories in which phasic dopamine encodes the perceived salience (*20*) or meaningfulness of events (ANCCR theory, (*19*, *26*, *74*)), and with reports that a subset of dopamine cells increases activity to a range of motivationally relevant events (*75*). Such a “valence-free” signal may be useful for adjusting learning rate, and/or for immediate control of attentional orienting and related processes (*57*, *76*, *77*).

In anterior Core, opposite net phasic dopamine responses to rewards and punishments are consistent with a valence-related signal, but there was little indication of RPE-like transfer to a reward-predictive cue. Higher dopamine tone and greater pharmacological response suggest that anterior Core is especially important for sustained affective/motivational states. The NAc is a key component of frontostriatal networks that maintain motivation to obtain rewards, and show hypoactivity in human depression (*78*). A more spatially nuanced view of NAc information processing may thus help understanding of symptoms such as anhedonia (*79*, *80*), and improve targeting for neuromodulation treatment of mood disorders (*81*).

Dopamine heterogeneity within the NAc has been most extensively examined in the coronal plane (especially Core vs medial Shell). Some investigations have found functional differences along the NAc anterior-posterior axis, primarily within Shell (*82–84*), but this axis has been underexplored for Core (*44*, *85*). Our discovery that there are at least two functionally distinct sets of Core dopamine signals may help explain variable results across prior studies—e.g. regarding aversive responses. Even when optic fibers are accurately placed within the Core atlas boundaries, photometry will sample variable proportions of these distinct signals (*86*). Furthermore, the precise anatomical locations of Core functional zones may vary between individuals, and even contribute to different motivational traits. Notably, in rats that tend to approach reward-predictive cues (“sign-trackers”; (*87*)) there is substantial transfer of Core dopamine reward responses to cue presentation (as seen here for posterior Core). In rats that instead approach reward delivery locations (“goal-trackers”) dopamine stays more anchored to reward (better resembling the anterior Core pattern). The extent of functional NAc territories can also be altered by an individual animal’s prior experience - e.g. stress exposure expands the volume of NAc Shell from which fearful behavior can be evoked (*83*).

Distinct dopamine dynamics within Core may reflect—at least in part—inputs from distinct dopamine cell subpopulations (*88–90*). Using detailed axon tracing, an example has been reported of a single VTA dopamine neuron that innervates solely anterior Core (*91*). Projections from genetically defined dopamine cell subpopulations known to innervate Core (e.g. *Sox6+, Cck+, Crhr1+*) appear not to be strongly selective along the anterior-posterior axis, though *Sox6+* innervation density may be higher in anterior compared to posterior Core (*92*, *93*). Differences in extracellular NAc dopamine may also be produced by local factors, such as variation in dopamine reuptake (*94*, *95*) and cholinergic modulation of dopamine axon activity (*96–100*). Downstream of dopamine, Core neurons show graded transcriptional differences along the anterior-posterior axis (*44*). The *Calcr+* subclass of D2R expressing cells are enriched in the posterior Core and project strongly to midbrain targets including VTA and midbrain raphe nuclei (*101*), a potential clue for deciphering the role of posterior Core dopamine.

In recent years the Core has been the principal testing ground for assessing computational theories of dopamine functions. These theories are now challenged by our demonstration that even just within the Core there are gradients of divergent dopamine signals. Of particular interest will be understanding how recipient Core neurons can make use of intermixed, rather than wholly separated, dopamine signals for both plasticity and online motivational control.

## Methods

### Animals

All experiments and procedures were performed in accordance with guidelines from the National Institutes of Health (NIH) *Guide for the Care and Use of Laboratory Animals* and approved by the UCSF Institutional Animal Care and Use Committee. Three wildtype C57BL/6J mice (RRID:IMSR_JAX:000664) and two DAT-IRES-Cre (B6.SJL-*Slc6a3^tm1.1(cre)Bkmn^*/J; RRID:IMSR_JAX:006660; adult (10–16 weeks at time of surgery) mice of both sexes (C57: 1M [Mouse 1], 2F [Mouse 2, Mouse 3]; DAT: 1M [Mouse 4], and 1F [Mouse 5]) were used for experiments. Mice were housed on a reverse 12-h light–dark cycle with lights off from 8:00 to 20:00, and all imaging was performed during the dark cycle; the holding room was maintained at ≈23–24 °C with 40–50% humidity. Throughout all experimental procedures, mice were given *ad libitum* access to food but were water-deprived to ∼85–90% of their pre-deprivation body weight, with daily weighing and adjustment of water allotment throughout conditioning; mice were not water-deprived for aversive-stimulation experiments. Sexes were pooled for all analyses.

### Surgery

Surgery was performed under aseptic conditions. Mice were anesthetized with isoflurane (5% induction, ≈1–2% maintenance), placed in a stereotaxic device (Kopf Instruments) and kept warm with a heating pad. Before incision, mice were administered carprofen (5 mg kg^−1^, subcutaneously (s.c.)) for pain relief, saline (0.3 ml, s.c.) to prevent dehydration and local lidocaine (1 mg kg^−1^, s.c.) to the scalp. After placing skull screws, a rectangular craniotomy was made on the right hemisphere from AP +0.25 to +2.25 and ML +1.1 to +3.0 relative to bregma. To record dopamine dynamics, AAV-DJ-CAG-dLight3.8 (Mouse 2, Mouse 3, Mouse 4, Mouse 5; final titer: 4 × 10^12^ vg ml^−1^) or AAVDJ-hSyn-dLight3.8 (Mouse 1; final titer: 8 × 10^12^ vg ml^−1^; Neurotools Viral Vector Core, University of North Carolina Chapel Hill) was injected into the nucleus accumbens (NAc). Virus was delivered at four sites (500 nL per site; 2 µL total) in a plane perpendicular to the medial-lateral axis at two depths along the DV axis at each of two sites along the AP axis (AP: +1.0 and +1.5; ML: +1.2-1.3; DV: −4.5 and −4.0 mm relative to bregma), through a glass pipette with a Nanoject III (Drummond Scientific) at a rate of 1 nL s^−1^. The pipette was held in place for 5–10 min after each injection to allow diffusion and then slowly retracted to prevent backflow up the injection tract. A 1.5 × 1.5 × 8 mm right angle microprism (*40*) (OptoSigma) was then implanted lateral to the injection sites to enable two-photon imaging of the NAc Core in the sagittal plane. A lateral, sagittal implant better preserves inputs from midbrain and prefrontal cortex compared to a coronal implant. The posterior, medial corner of the prism was implanted at AP: +0.5, ML: +1.4-1.5, DV: −5.0 mm relative to bregma. The prism was held with a micro bulldog clamp (WPI 14119) covered in heat shrink tubing and attached to the stereotaxic frame via a cannula holder attachment (KOPF 1766-AP). After removing the dura, the prism was quickly lowered into the brain to about −2.5 mm DV, and then lowered in steps of 0.2 mm every 90-120 s. When the prism reached −3.5 mm DV, it was then lowered in 0.1 mm steps every 90-120 s until reaching −5.0 mm. The prism was then affixed to the skull with C&B Metabond (Parkell). A custom-designed stainless steel head ring (5-mm inner diameter, 11-mm outer diameter, 3-mm height) for head fixation was then placed over and around the prism and secured to the skull with cyanoacrylate glue. Finally, the head-ring, prism, and skull screws were covered in dental cement to anchor the prism and head-ring to the skull. The DAT-IRES-Cre subset of mice (Mouse 4 and Mouse 5) received an additional virus injection for mScarlet expression (AAV5-Ef1alpha-mScarlet; Addgene 131002, final titer: 1 × 10^13^ vg mL^−1^) in VTA dopamine neurons (AP: −3.1, ML: +0.5, DV: −4.4 mm relative to bregma). All virus injections and prism implants were in the right hemisphere. Following surgery, mice were given buprenorphine (0.1 mg kg^−1^, s.c.) for pain relief and allowed at least 8 weeks for recovery and viral expression before imaging.

### Two-photon imaging

dLight imaging was performed using an Ultima2pPlus resonant-scanning two-photon microscope (Bruker) coupled to a MaiTai Deepsee Ti:Sapphire tunable ultrafast laser (Spectra-Physics), and a long-working-distance air objective suited to microprism-coupled imaging (Cousa objective (*102*); ×10, NA 0.50, 20-mm working distance). In the subset of animals with mScarlet expression in their dopamine axons (**Fig. 1G**), mScarlet was additionally imaged at 1050 nm with a FemtoFiber Ultra 1050 laser (Toptica). Average power at the objective was kept below 100 mW at all imaging wavelengths. Standard acquisition consisted of 64 × 64 pixel recordings (22.1 µm px^−1^) at ≈205 Hz. For high-resolution imaging, 512 × 512 pixel recordings (2.76 µm px^−1^) were acquired at ≈30 Hz. Imaging frames, behavioral events, stimulus times and camera frames were synchronized through a shared voltage record. Raw frames were motion-corrected by rigid registration in Suite2p (*103*).

#### Unpredicted random reward

Sucrose (15% w/v, ≈3 µl/drop) was delivered though a lick spout at exponentially distributed inter-reward intervals (IRIs), constrained to 12–18 s or 60–90 s, with mean IRIs of 15 or 75 s, respectively. For reward-response maps and ROI PSTHs, 128 rewards per mouse were pooled (64 rewards per IRI condition, **Fig 3H-J**); wavelength-characterization (**Fig 1C-E**) and multi-scale ROI analyses (**Fig 1H**) used 15-s IRI sessions (32 rewards), and pharmacology sessions used 75-s IRI (**Fig 1F**, **Fig 2D-F**).

#### Pavlovian (cue-reward) conditioning

A CS+ (12 kHz pure tone, 250 ms) predicted sucrose after a 2 s trace interval following cue offset (reward delivered 2.25 s after cue onset; 100% reinforced), and a CS− (3 kHz, 250 ms) predicted no reward; cues were randomly interleaved (50 trials per cue, 30 s inter-trial interval). These tones have been previously shown to produce reliable anticipatory licking (*26*), and were thus not counterbalanced across animals. Cue, reward (solenoid), and lick events were controlled and recorded with an Arduino-based behavioral controller running a custom MATLAB graphical user interface (B-CALM)(*104*). Licks were detected by contact between the mouse’s tongue and the lick spout completing an electrical circuit; each lick was timestamped on the Arduino together with cue and reward events and logged in MATLAB for offline analysis. Our main measure of learning in this task was anticipatory licking (between cue onset and reward delivery). All but one mouse (Mouse 1) were imaged on the first day, and after 14 conditioning sessions, imaging was performed every three sessions (15, 18, 21, and 24). The first imaging session in which the CS+ lick rate exceeded 1.40 Hz and the CS− lick rate was below 0.70 Hz was designated the “final” conditioning session. One mouse (Mouse 2) failed to learn (CS+ lick rate remained below 0.5 Hz at session 24) so was excluded from analyses.

#### Aversive stimulation

Noxious heat was delivered to the whisker pad using a 1450 nm infrared laser (Edmund Optics #70-230; 200 ms pulses; 60 s inter-stimulus interval; 30 stimulations per session) coupled through a patch cord and collimator. The laser output was approximately 450 mW at the patch-cord tip, and the beam diameter at the whisker pad was approximately 3 mm. The pulse duration was set to 200 ms, which reliably evoked an orofacial reaction (“face-wipe”) (*61*). To quantify face-wipe events, the face and front paws were filmed at 50 fps (Allied Vision #14184) and the paw and mouth were tracked with Facemap (*105*) (v1.0.8). Face-wipe events were identified when the paw y-position reached 80% of the distance toward the median mouth y-position across the recording. Wiping probability was calculated as the fraction of frames classified as wipe events and was averaged across stimuli. Aversive stimuli were not given to one mouse (Mouse 1).

#### Excitation-wavelength characterization

In separate sessions, we excited at 920 and 820 nm during delivery of unpredictable sucrose rewards (mean inter-reward interval: 15 s), and in a subset of mice we swept the excitation from 760 to 960 nm in 20 nm steps. To counterbalance the order of wavelengths, wavelengths were swept in both directions, and reward responses were measured 10 times at each wavelength in each sweep direction, yielding a total of 20 reward responses per wavelength.

#### Pharmacology

Cocaine hydrochloride (10 mg kg^−1^; Sigma-Aldrich, C5775), SCH-23390 hydrochloride (0.3 mg kg^−1^; Tocris Bioscience, 0925) or vehicle (sterile 0.9% saline, 10 mL kg^−1^ body weight) were administered on separate sessions in a counterbalanced order across mice, with at least two days between sessions to allow for washout. During injection, the PMT was shuttered so that the room lights could be turned on to facilitate accurate subcutaneous injection while the mouse remained head-fixed.

#### Histology

After the conclusion of experiments, mice were transcardially perfused with 4% paraformaldehyde. Brains were then postfixed in 4% paraformaldehyde in the intact skull to maintain prism geometry during fixation. After postfixing, brains were carefully removed from the bottom of the skull and embedded in 5% low-melting point agarose for vibratome sectioning. Brains were sectioned horizontally at 50 µm. After sectioning, every other slice was stained for calbindin to differentiate Core from Shell. Note that we capitalize Core and Shell throughout the manuscript to better distinguish the brain regions from ordinary usage of these words. Slices were blocked using a PBS 1x (Sigma-Aldrich P4417) solution containing 1% Triton X-100 (Sigma-Aldrich, 93443), and then incubated with primary rabbit anti-Calbindin (D1I4Q, CST 1:500) antibodies with normal donkey serum (1:50). This was followed by a secondary donkey anti rabbit antibody conjugated with Setau-647 (AB_3095048, LifeCanvas 1:250) with normal donkey serum (1:50). Slices were then mounted with DAPI Flouromount-G. To examine dLight and mScarlet (when applicable) expression in addition to calbindin staining, slices were imaged using a Keyence BZ-X800 microscope with a 2x objective. Slices containing the anterior commissure, the clearest anatomical landmark for Core, and the most ventral slice containing a detectable imprint in the brain from the prism were reimaged with a 10x objective (**fig. S2**). Stitched images of full brain slices were then cropped to focus on either the hemisphere containing the prism (anterior commissure slice) or the striatum (most ventral slice).

#### Registration of imaging FOV to anatomical atlas

We first took z-stacks (512 x 512 resolution, 20 µm steps, each slice the average of 512 frames imaged at 30 Hz) through the full prism FOV from the prism surface through 260 µm more medially into the tissue (**fig. S1**). This allowed us to visualize the anterior commissure, the clearest anatomical landmark for Core, in all mice. Slices from these z-stacks containing the anterior commissure were then manually registered to the sagittal atlas (*106*) to estimate the dorsal-ventral and anterior posterior extent of the imaging FOV. These position estimates were later confirmed through histology (**fig. S2**).

#### Anatomical ROI definition

Using the determined anatomical placements of the prism FOV for each mouse, anterior and posterior ROIs were selected in regions of the anterior-dorsal Core and posterior-ventral Core that were present in the imaging FOV of all mice **(fig. S1).** As the FOVs between mice did not perfectly overlap, ROIs were placed differently across mouse’s FOV to match equivalent anatomical positions in the dorsal-ventral and anterior-posterior axes. When mapped onto the 64×64 pixel images, these ROIs corresponded to a 12 × 12-pixel square (≈265 × 265 µm) used for all ROI level comparisons. For the smaller ROIs used in spatial maps of the time course of event-evoked dopamine (**Fig. 5C, S9, S11**), time course traces acquired at ≈205 Hz were downsampled to ≈51 Hz (4× binning) then Savitzky-Golay filtered (2^nd^ order, ≈60 ms window).

### Imaging analysis

#### dLight ratio calculations

For analyses requiring correction for local sensor expression — tonic-level maps and pharmacology (**Fig. 2**), reward responses (**Fig. 3**), cue responses (**Fig. 4**), and aversive-stimulus responses (**Fig. 5**) — we used the “dLight ratio” obtained by normalizing the dopamine-sensitive 920 nm signal to a baseline fluorescence (F₀) derived from the isosbestic 820 nm signal. The 820 and 920 nm reference images were co-registered, and a single linear fit relating the two was used to scale and convert the 820 nm image into the baseline F₀. The dLight ratio was then computed as (F_920_ − F_820_)/F_820_, then applied to each pixel. After applying the estimated alignment shift to co-register the two images, pixels with low 920 nm fluorescence were excluded from the fit and from subsequent dLight ratio analysis (see Pixel selection, and unified masking below). This retained approximately 85% of the field of view on average.

#### Cross-session registration

Because the precise FOV drifted very slightly between sessions, the per-session translational shift was estimated by subpixel phase-correlation (Guizar-Sicairos DFT) to a common reference image and used to correct each ROI center, so that the same anatomical footprint was sampled across sessions.

#### Pixel selection, and unified masking

To restrict the analysis to a reliable fluorescence signal and exclude background or imaging artifacts, pixels were selected in two steps. First, the imaged field of view was delimited by excluding dark, out-of-prism regions: pixels below 15% of the 920 nm 99th-percentile intensity were removed. Second, within this field, a signal mask retaining dLight-expressing pixels was defined by Otsu’s method (*107*), which selects the intensity that best separates foreground from background by maximizing between-class variance of the histogram; the Otsu threshold was computed over field-of-view pixels only (so that out-of-prism background did not bias it) and then relaxed (×0.4) to keep weakly labeled pixels. To enforce an identical footprint across conditions, each session’s mask was aligned to a common reference, and a unified mask was taken as the intersection across sessions (i.e., a pixel was kept only if it passed threshold in every session).

#### Diagonal band analysis

A single diagonal band (a rectangle), from the posterior-ventral to the anterior-dorsal corner of the field of view, was defined identically for every mouse. The band was 15 pixels wide, and signals were averaged across the width at each position along the diagonal. This diagonal band was used to generate spontaneous-activity kymographs (**Fig. 3B**) and to quantify the spatial slope of aversive responses (**Fig. 5D**).

#### Spontaneous activity

dLight signals were recorded (≈205 Hz) for about 20 min in the absence of rewards or other stimuli and down-sampled to ≈40 Hz for subsequent analyses. For kymograph and spectral clustering analyses, dF/F was referenced to the per-pixel temporal mean of the 920 nm recording, detrended by subtracting a slowly varying baseline estimated as the 8th percentile within a centered 30 s sliding window, and lightly smoothed (Savitzky–Golay, ≈0.2 s window). Two anterior–posterior domains were identified per mouse by applying unsupervised spectral clustering to dF/F time courses from individual pixels within the signal mask, and selecting two clusters (MATLAB spectralcluster). Power spectra were estimated by Welch’s method (100 s Hann windows, 50% overlap), using data collected outside of a specific task.

#### Pharmacology analysis

For each session, a pre-injection map was averaged over the 10 min immediately preceding shutter closure, and a post-injection map was averaged over a fixed 10 min window spanning 30–40 min after shutter closure. Sucrose rewards were delivered at 75 s inter-reward intervals on average during these periods, and frames within 10 s after each reward were excluded from both windows to minimize the phasic reward-related component. Post-injection maps were rigidly registered to the pre-injection map.

### Fluorescence lifetime imaging (FLIM)

FLIM was performed on a two-photon microscope using time-correlated single-photon counting (TCSPC; Bruker TT-FLIM2 module). Photon arrival times were recorded over a 12.5 ns window, matching the 80 MHz laser repetition period, with 100 mW excitation power, 30 µs pixel dwell time, and 128 x 128 pixel acquisition. Each FLIM recording lasted 1,400 s, during which mice received uncued rewards at a mean inter-reward interval of 75 s. To isolate baseline activity, frames within 5 s following each reward were excluded, and photons from the remaining frames across the two sessions per mouse were pooled and spatially binned 2 × 2 before lifetime estimation. Lifetimes were estimated using a phasor approach. The instrument response was measured from second-harmonic generation (SHG) of mouse-tail collagen, an effectively zero-lifetime signal, acquired under settings identical to the dLight imaging (*108*). TCSPC decays were transformed into phasor coordinates by discrete Fourier transform at the fundamental modulation frequency (80 MHz) and referenced to the spatially integrated SHG phasor by complex division to correct for instrument delay and demodulation. The phase lifetime was calculated as τφ = tan(φ)/ω, where φ = atan2(s, g) (*109*, *110*). Calibration was validated using Coumarin 6 (τφ ≈2.46 ns; expected ∼2.4–2.5 ns). Longer τφ therefore indicates a greater fraction of dopamine-bound sensor molecules.

### Statistics

No statistical test was used to predetermine sample sizes. All statistical tests were two-tailed. Comparisons between anterior and posterior ROIs were made on scalar per-mouse summary values using two-tailed paired t-tests (drug-induced dLight ratio, slow- and fast-band PSD power during spontaneous activity, cue-, reward-, and aversion-evoked dLight ratios); comparisons of a single response against zero used two-tailed one-sample t-tests (drug-induced dF/F, unpredicted reward-, cue- and predicted reward-evoked dLight ratios, spatial slope of aversion response). n denotes the number of mice and is stated for each comparison (n = 5 for tonic, pharmacology and reward; n = 4 for cue-reward and aversion), with each mouse contributing one value per condition. Data are mean ± s.e.m. Results were considered significant at α = 0.05 (*p < 0.05, **p < 0.01, ***p < 0.001; n.s., p > 0.05). All analyses were performed in MATLAB R2024a (MathWorks).

## Acknowledgments

We thank the following for careful reading of prior versions of this manuscript: Veronica Alvarez, Kevin Bender, Robert Edwards, Arif Hamid, Elyssa Margolis, Khaled Moussawi, Matthew Pomrenze, Tommaso Patriarchi, Bernardo Sabatini and members of the Berke and Namboodiri laboratories. **Funding:** This work was supported by the National Institutes of Health (R01DA045783 (J.B.); R01MH129582 (V.M.K.N.); F32DA060044 (D.A.B.)), Alfred P Sloan Fellowship (V.M.K.N.), Pew Biomedical Scholarship (V.M.K.N.), Klingenstein-Simons Fellowship (V.M.K.N.), the Scott Alan Myers Endowed Professorship (V.M.K.N.), the University of California, San Francisco, and the State of California. Author contributions: Conceptualization: V.M.K.N, J.B.; Methodology: S.K., D.A.B., C.K., H.J.; Investigation: S.K., D.A.B., C.K. F.F.; Funding acquisition: V.M.K.N, J.B.; Supervision: V.M.K.N, J.B.; Writing – original draft: S.K., D.A.B., V.M.K.N, J.B; Writing – review & editing: S.K., D.A.B., V.M.K.N, J.B. **Competing interests:** The authors declare that they have no competing interests.

---

**Figure S1:**
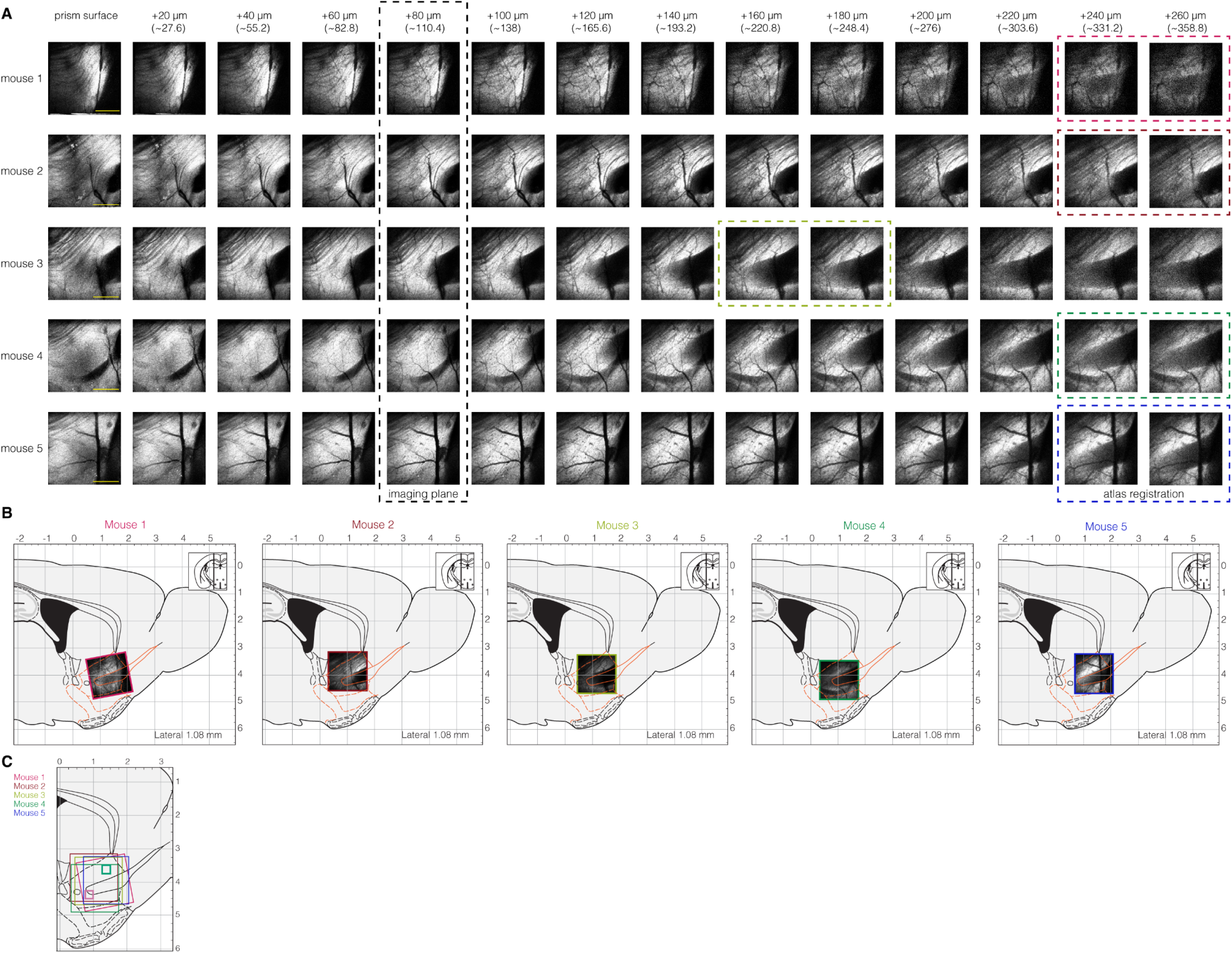
Anatomical registration. A. Z-stacks of full FOV (Mouse 1: 1.509 x 1.509 mm, all other mice: 1.415 × 1.415 mm) from prism surface to (nominal) 260 µm deeper (more medial) into the tissue. Numbers listed on top are the distances moved by the microscope for each slice relative to the prism surface, while numbers in parentheses represent the calculated depth moved through the brain tissue (assuming a refractive index of 1.38 for striatal tissue, and not correcting for non-paraxial effects). Each image is the mean of 512 frames with 920 nm excitation and with a resolution of 512×512 pixels. Black dashed rectangle marks the imaging planes for each mouse used for experiments throughout the study (chosen based on maximum average signal intensity). Colored dashed lines represent sections averaged for atlas registration in B. Note that individual image slices are auto-scaled from the 5th to the 95th percentile of pixel intensities to maximize contrast for display. Scale bars: 500 µm. B. Images from z-stacks (**A**) containing the anterior commissure (the clearest anatomical landmark in the FOV) manually registered to the sagittal atlas (Franklin & Paxinos, 2008) to determine the dorsal-ventral extent of the imaging FOV. Anterior-posterior position estimate was confirmed through histology (**fig. S2**). FOV image shown on atlas section represents the average of the two slices outlined with colored dashed boxes in A. Numbers on sides of atlas section represent distance from bregma in mm. C. Atlas section overlaid with outlines of full FOV for all five mice. Green and pink squares represent anatomically consistent anterior and posterior ROIs, respectively, used throughout the study (see Methods).

**Figure S2:**
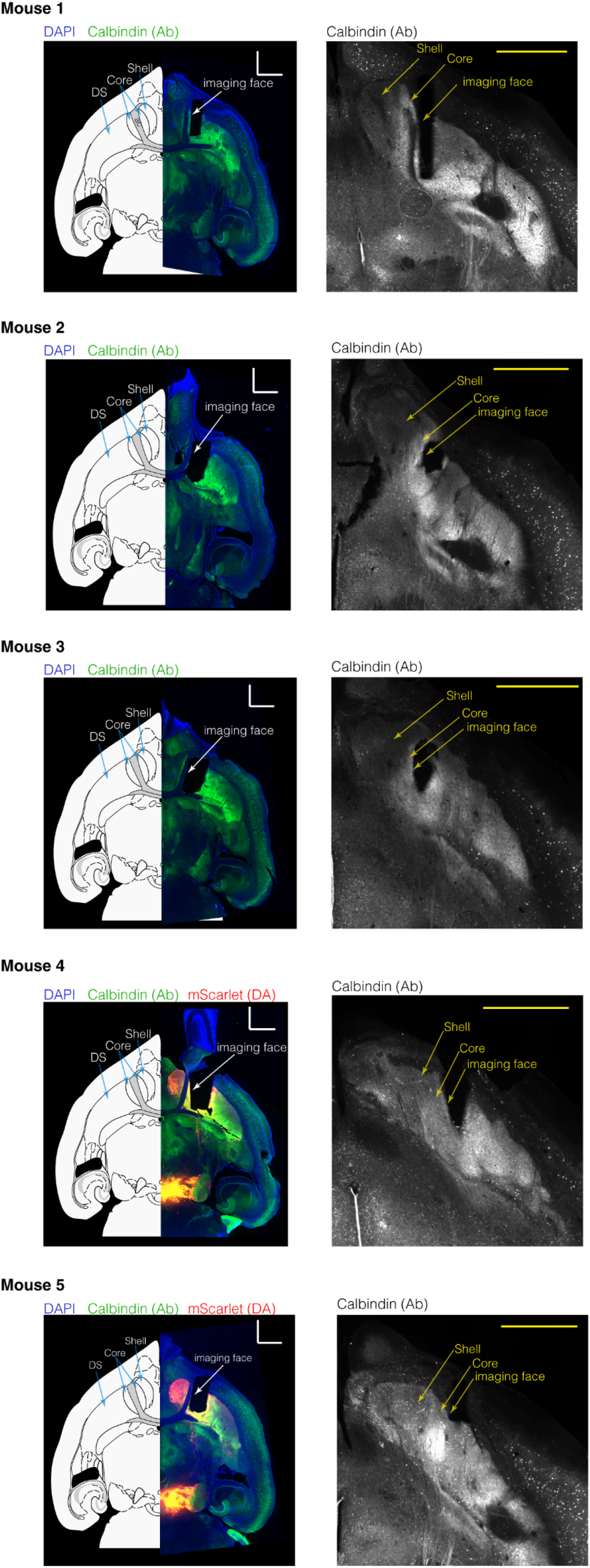
Histology confirms prism targeting of Core. *Left,* Horizontal brain sections (50 µm) containing the anterior commissure for visual landmarking aligned to the best matched horizontal atlas section (Franklin & Paxinos, 2008) (−4.12 mm ventral to bregma). This represents roughly the middle of the prism imaging area in the dorsal-ventral axis. Consistent with *in vivo* imaging (**fig. S1**), prism implants in all mice were lateral to the anterior commissure (gray structure bisecting Core on atlas diagram). Sections were stained with DAPI (blue), and antibody against calbindin (green), a standard marker for distinguishing accumbens Core from the calbindin-negative Shell. Sections were imaged with a 10x objective on a Keyence BZ1000 slide scanning microscope and stitched. For Mouse 4 and Mouse 5 (both DAT-IRES-Cre), a DIO-mScarlet virus was injected into the VTA to label dopamine neurons and axons (red). As calbindin was imaged in the far-red channel, there is some bleed-through from mScarlet in those mice. DS: dorsal striatum. Scale bars: 1 mm. *Right,* Closer views of calbindin staining, using the most ventral section in each brain with a detectable mark from the prism. (The ventral-most section is chosen as this is most likely to include Shell, if present at all). Core and Shell arrows point to higher and lower areas of calbindin expression in the region of the nucleus accumbens.

**Figure S3:**
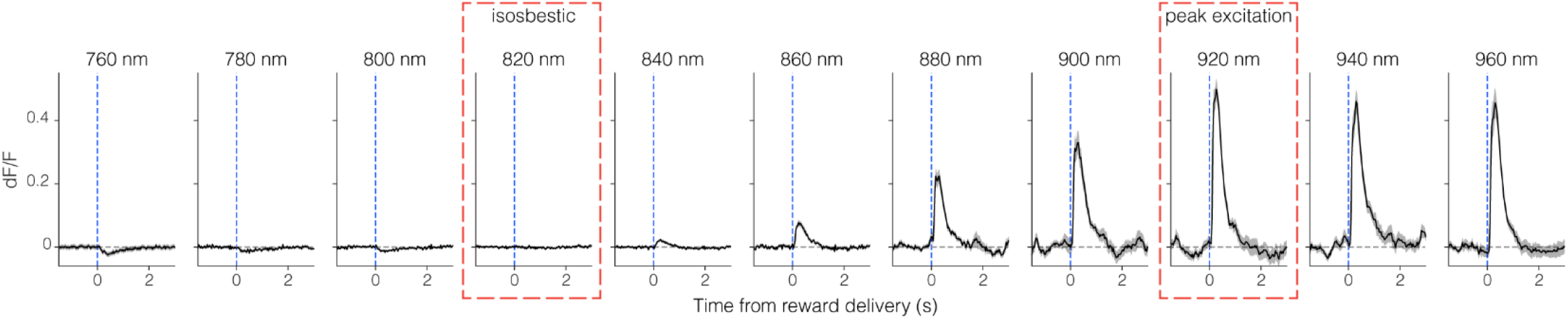
dLight3.8 signal response to reward depends on excitation wavelength. Averaged dLight time courses centered on unpredictable reward delivery, for a range of two-photon excitation wavelengths in a single animal (Mouse 1). N=20 reward deliveries at each wavelength: 10 rewards at each wavelength while progressively sweeping from 760nm to 960nm, then another 10 rewards each while reversing the order (960nm to 760nm).

**Figure S4:**
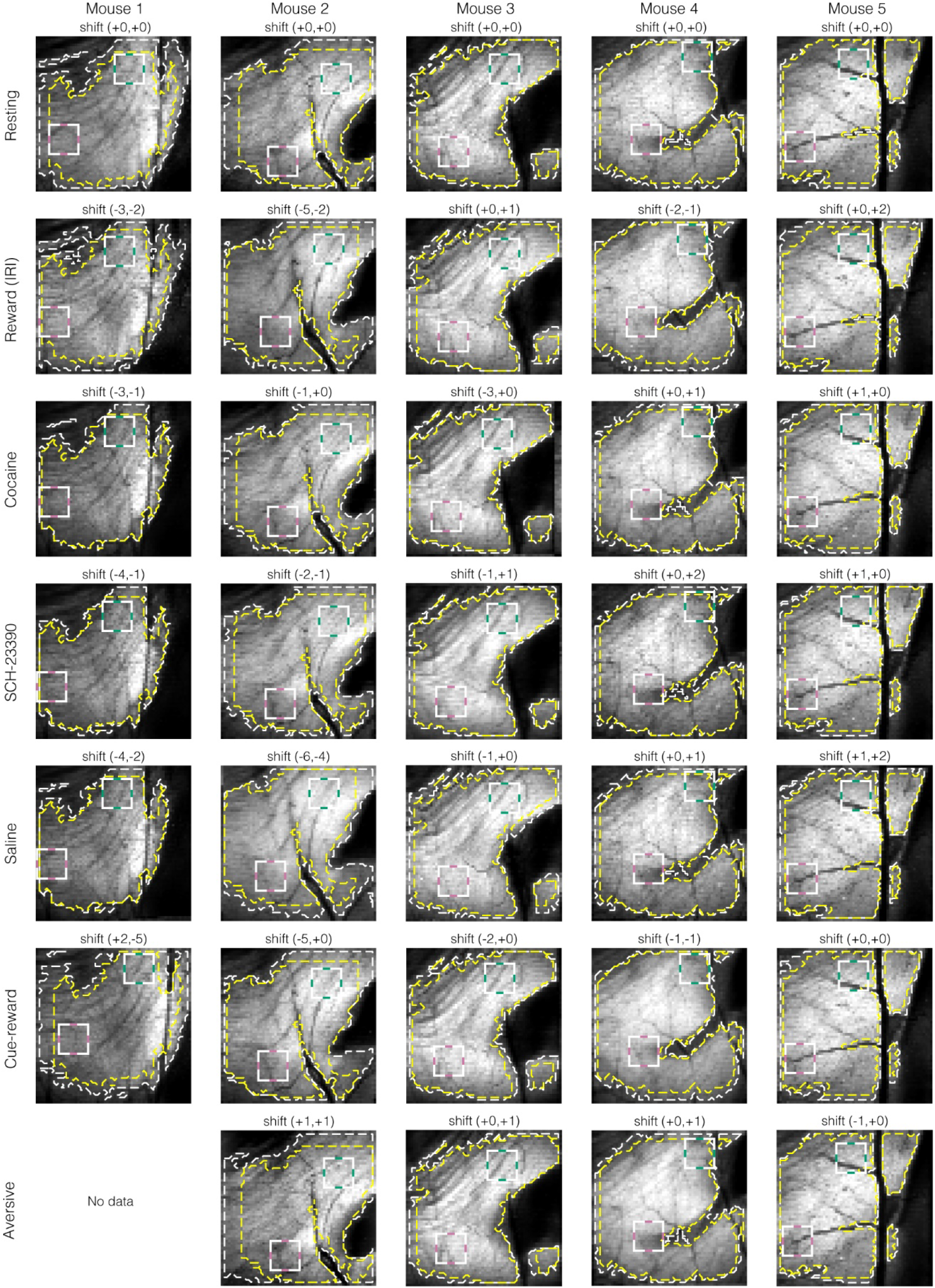
Signal masking and cross-session alignment of fields of view. To ensure a consistent FOV was analyzed across experiments, and that low-signal pixels did not bias analyses, each session’s FOV was registered to a common reference based on mean images acquired during rest periods. After alignment, FOVs were threshold masked to exclude low-signal pixels (see Methods). For each mouse (columns) and experiment/session (rows), the mean fluorescence images (grayscale) are shown with the signal masks overlaid. White dashed contours outline each session’s own signal mask, and yellow dashed contour outline the intersection of masks across sessions. These intersection masks were used for all analyses: i.e. a pixel was included only if it passed the masking criterion in every session. Anterior (green/white) and posterior (pink/white) ROIs are shown for reference. Labels above each image indicate the spatial shift (in pixels) applied for alignment.

**Figure S5.**
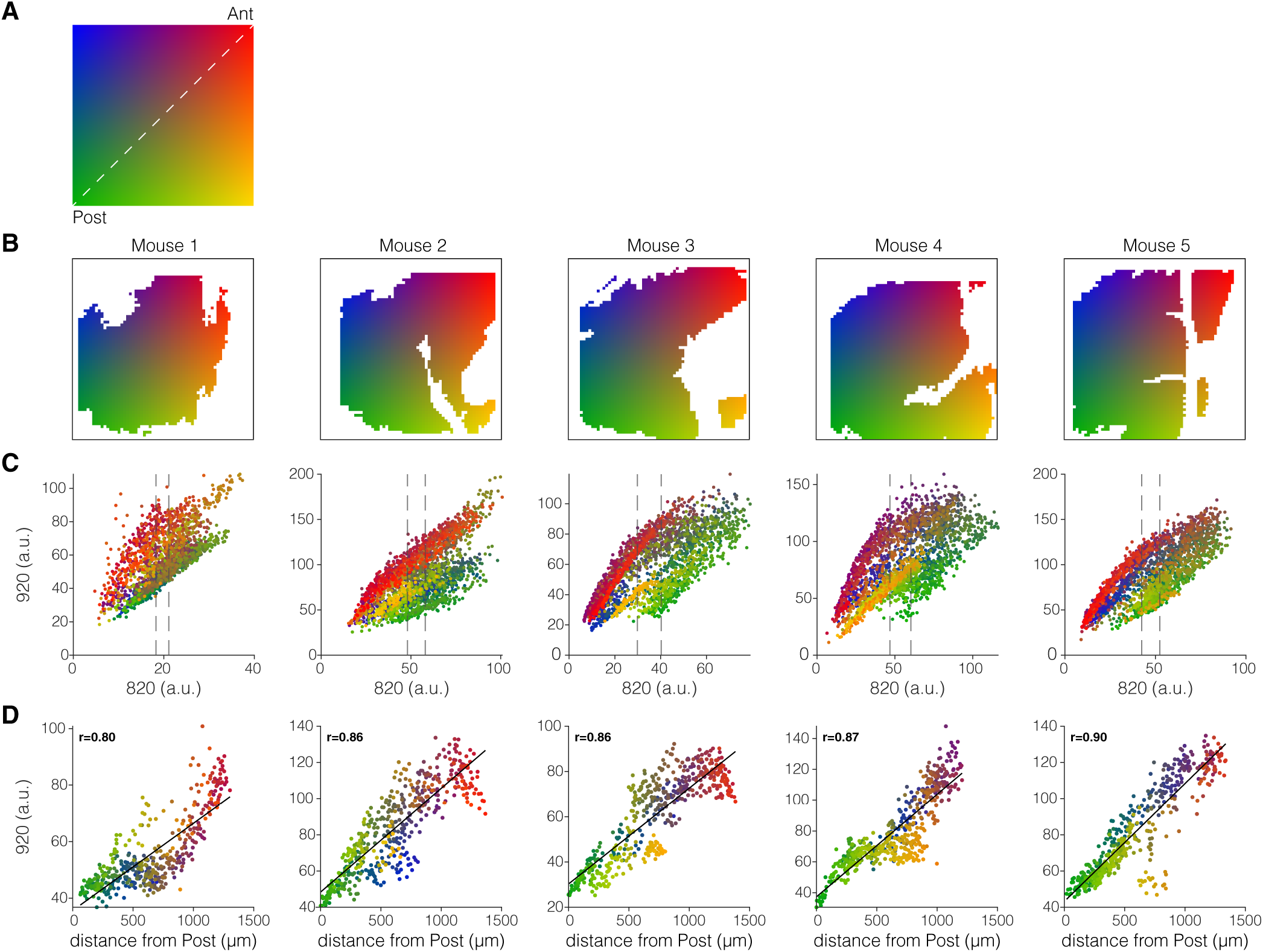
Anterior-posterior dopamine difference is not the result of dLight expression. **A)** Position color code within the field of view (FOV). The posterior/ventral (“Post”)-to-anterior/dorsal (“Ant”) axis runs from green to red. **(B)** Position color code applied to pixels from five mice. White regions indicate excluded pixels. **(C)** 920-nm versus 820-nm fluorescence for each pixel, colored by position. Dashed lines indicate the 40th and 60th percentiles of the 820-nm intensity distribution used in (D). **(D)** 920-nm fluorescence versus position along the posterior/ventral-to-anterior/dorsal axis for pixels within the 40–60th percentile 820-nm band. Black lines show linear fits. Insets show Pearson’s r (Mouse 1: p = 4.1e-97, n = 436 pixels; Mouse 2: p = 4.6e-142, n = 436 pixels; Mouse 3: p = 6.9e-144; n = 491 pixels; Mouse 4: p = 3.9e-163, n = 528 pixels; Mouse 5: p = 2.8e-183, n = 497 pixels). The 920-nm signal increased along the axis in all five mice when controlling for similar 820-nm intensity.

**Figure S6:**
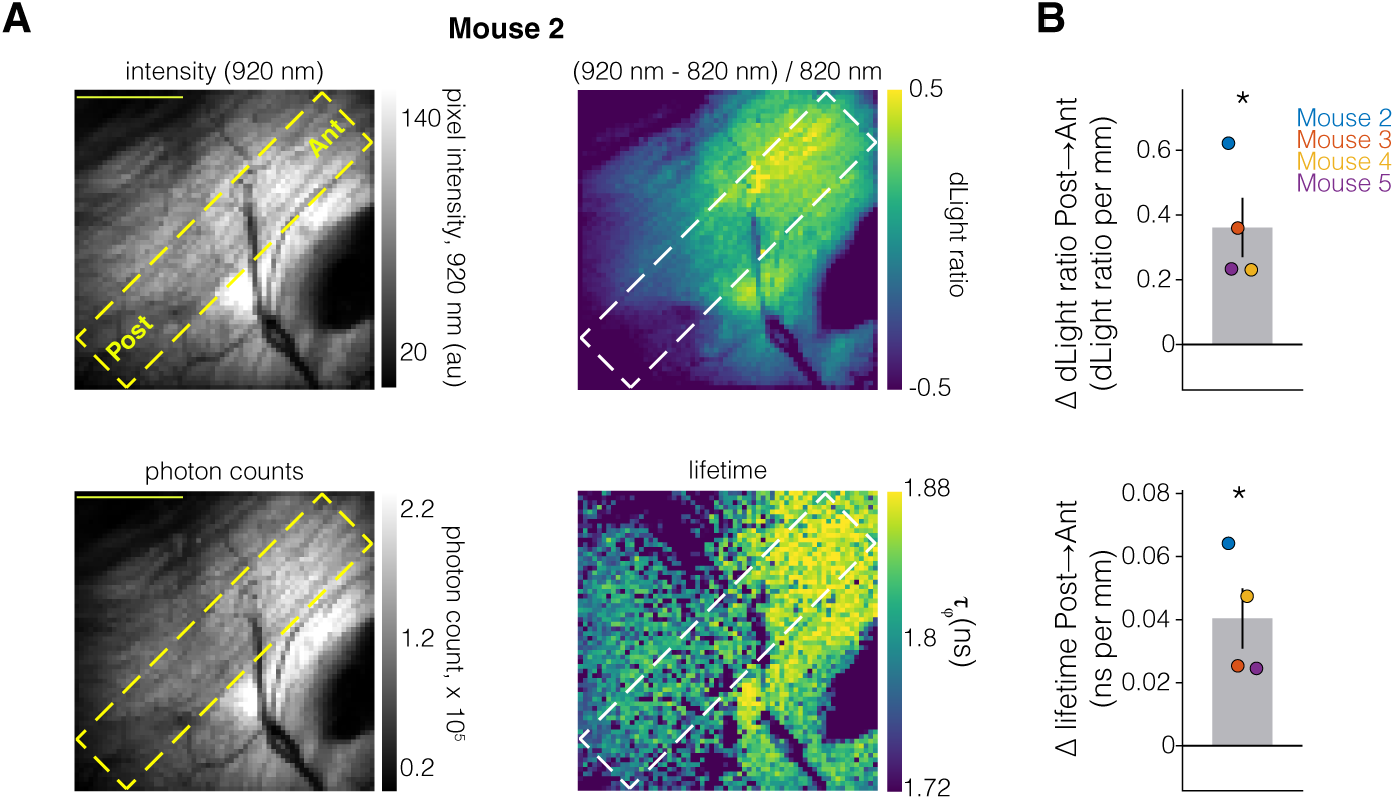
Fluorescence lifetime imaging confirms greater tonic dopamine in anterior Core. **A.** Field of view from example mouse (Mouse 2). *Top left*, mean fluorescence image at 920 nm; yellow dashed box indicates diagonal ROI stretching from posterior (Post) to anterior (Ant). *Top right*, corresponding dLight ratio map, calculated as (920nm – fitted 820nm)/fitted 820nm. *Bottom left*, accumulated photon-count image for lifetime estimation. *Bottom right*, phasor-derived fluorescence lifetime (τφ) map. Longer τφ indicates a greater dopamine-bound fraction (see Methods). Maps show pixels passing the signal mask. Scale bar: 500 µm. **B.** Posterior-to-anterior slopes of the dLight ratio (top) and fluorescence lifetime (bottom) (n = 4, Mice 2,3,4,5). Positive values indicate an increase toward anterior Core. Bars, mean ± SEM; points, individual mice. One-sample t-test vs. zero slope: dLight ratio: t(3) = 3.93, p = 0.0293; lifetime: t(3) = 4.23, p = 0.0242.

**Figure S7:**
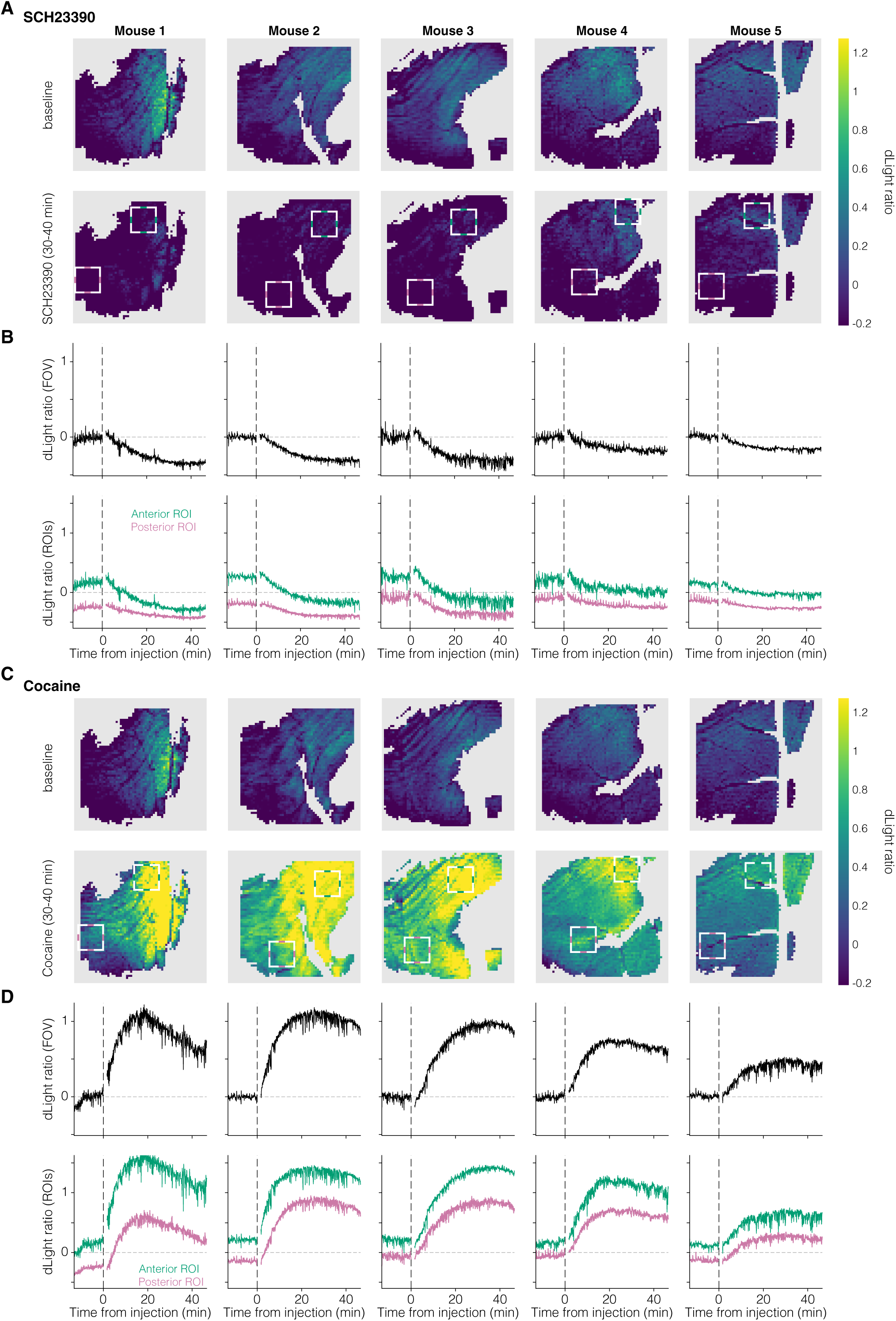
Maps of dopamine responses to drugs, without baseline subtraction. **A.** Mean dLight ratio maps before (top) and after (bottom) SCH-23390 injection for each mouse. Baseline: 10 min before injection; post-injection: 30–40 min later. White squares indicate anterior (upper right) and posterior (lower left) ROIs. Gray, pixels outside the analysis mask. **B.** dLight-ratio time courses for each mouse. Top, field-wide average; bottom, anterior (green) and posterior (pink) ROI averages. Traces are shown in 5-s bins; dashed line, injection. Data were not recorded during the injection itself. **C-D.** As A, B, for cocaine.

**Figure S8:**
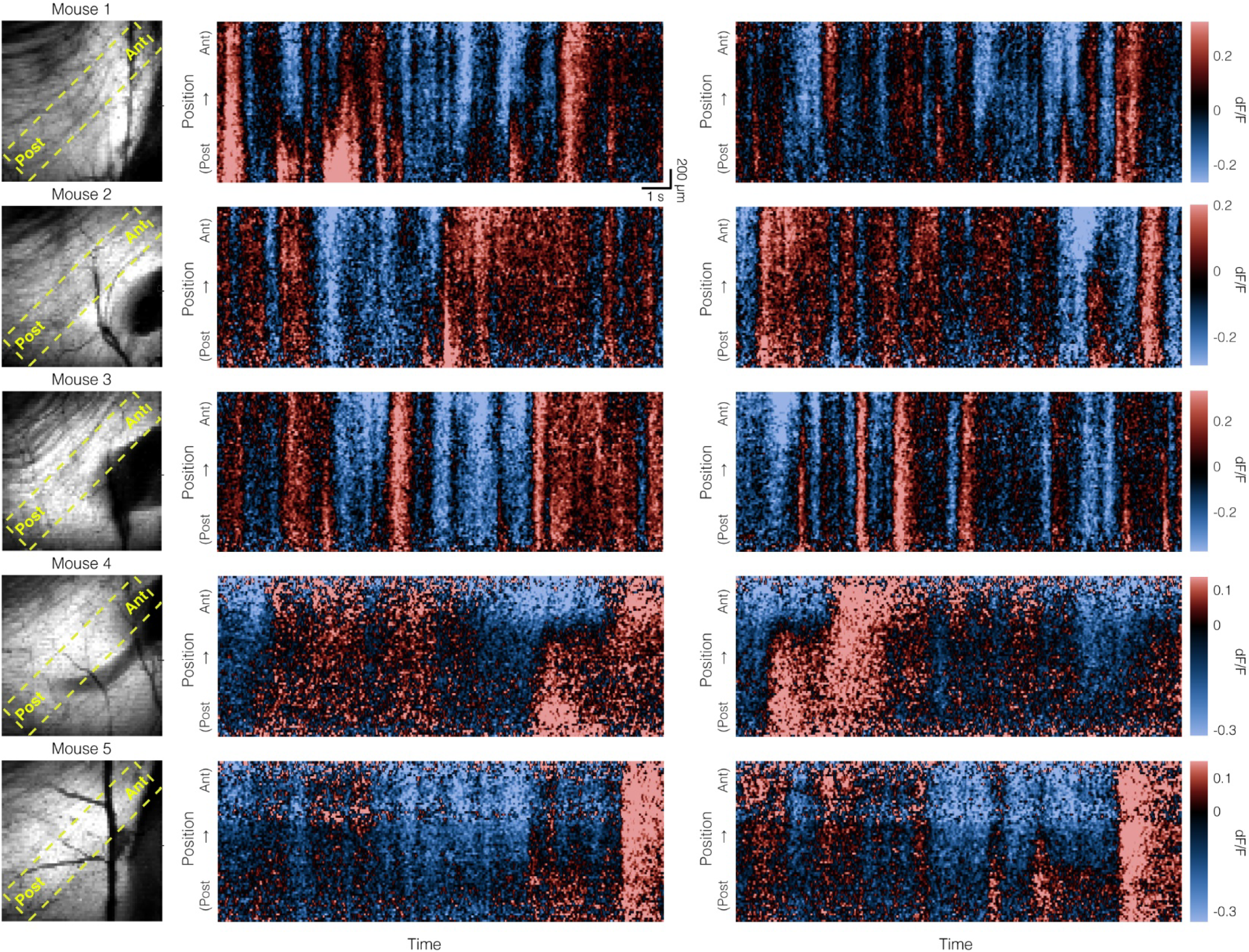
More examples of spatially heterogeneous spontaneous activity within Core. Data from each of the five mice, in the same format as Fig. 3A-B.

**Figure S9:**
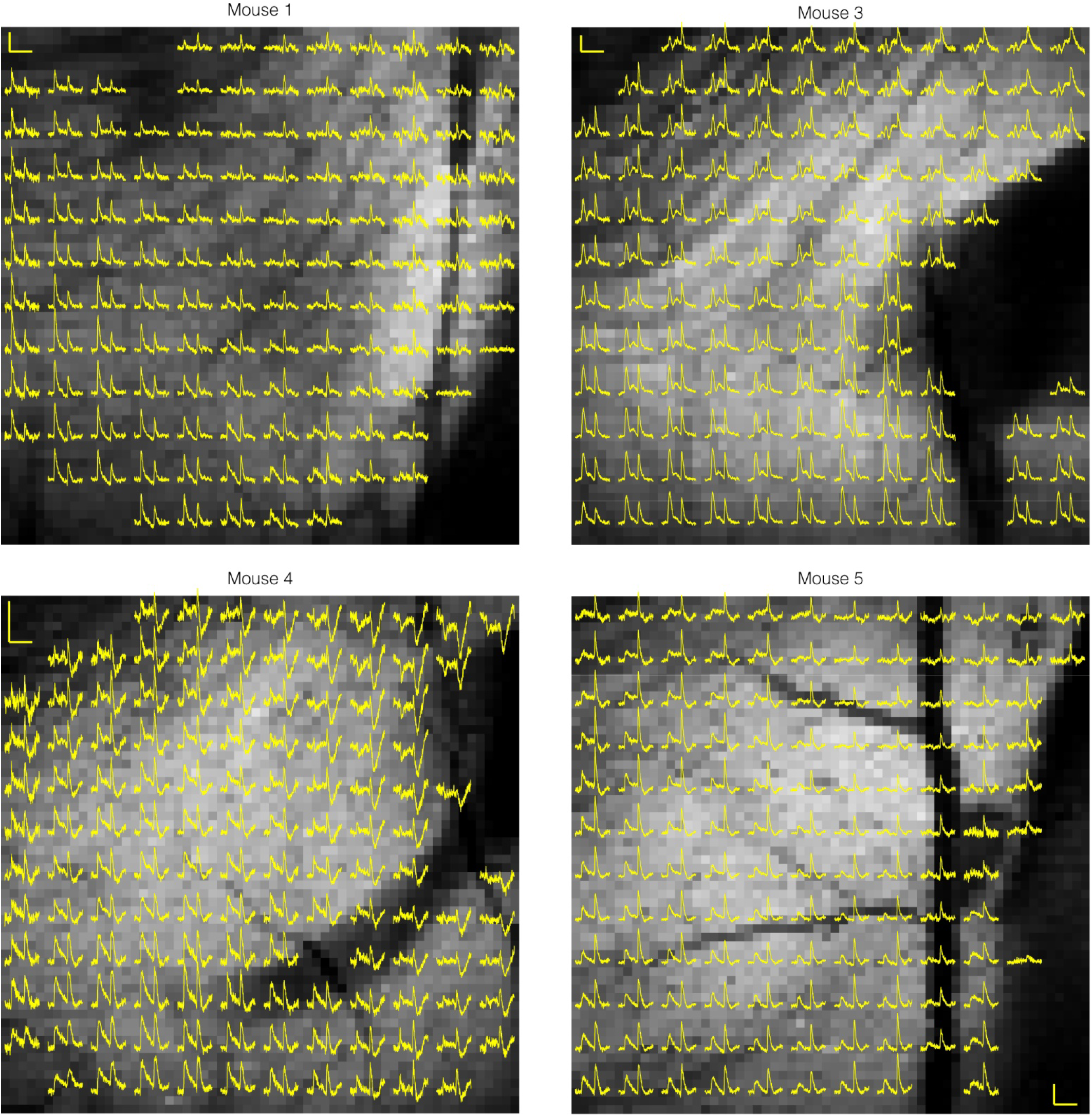
Cue and reward responses across Core in individual mice after conditioning. For each mouse, the full FOV (greyscale) was divided into a grid of 5 × 5 pixel zones (≈22 µm/pixel, recorded at ≈205Hz). Yellow traces show dLight times averaged within each zone (−1 – 5s relative to CS+ onset; 50 CS+ events) on the final day of conditioning (mean of 50 CS+ trials stimulations, from −1 – 5 s relative to CS+ onset). Scale bars: 0.3 dLight ratio, 4 s.

**Figure S10:**
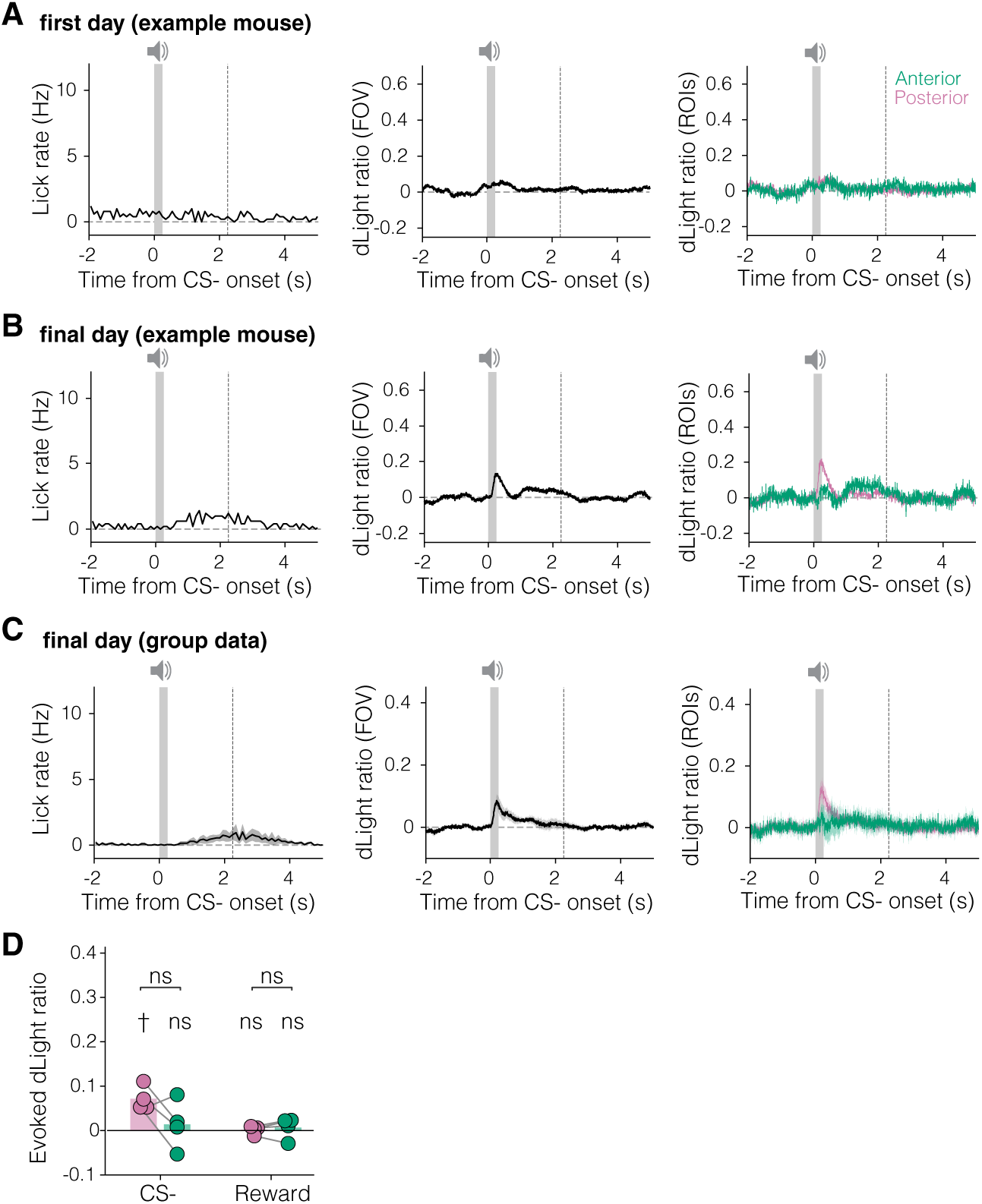
CS-responses during cue-reward conditioning. **A-C** Same as Fig. 4 A,C,E, but for CS-cues. **D**. Same as Fig. 4F, but for CS-cues. Quantification of dopamine responses to CS− (0–0.5 s) and during the 2.25–2.75 s window after CS− onset, corresponding to the reward-response window on CS+ trials, in anterior (green) and posterior (pink) ROIs. One-sample t-tests (n = 4 mice) comparing to zero: posterior CS−, t(3) = 5.24, p = 0.014; anterior CS−, t(3) = 0.501, p = 0.651; posterior 2.25–2.75 s, t(3) = 0.366, p = 0.739; anterior 2.25–2.75 s, t(3) = 0.549, p = 0.621. Anterior and posterior ROIs were also compared to each other using paired t-tests: CS−, t(3) = 1.926, p = 0.15; 2.25–2.75 s, t(3) = −0.632, p = 0.572.

**Figure S11:**
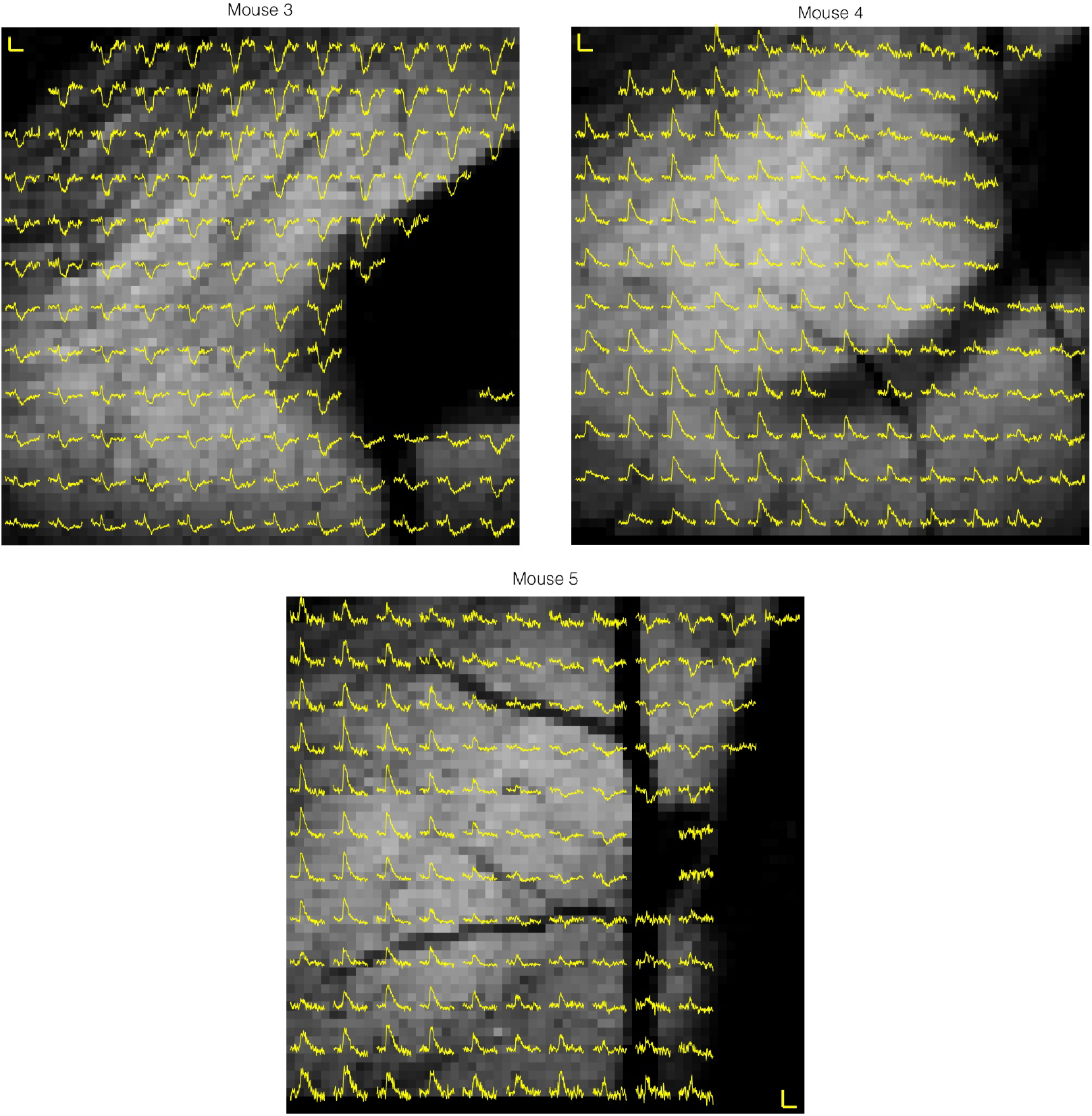
Aversive stimulation response maps for additional mice. Same format as Fig. 5C, for the other mice. Here the underlying greyscale images are at 64×64 resolution. Scale bars: 0.2 dLight ratio, 1 s.

**Figure S12:**
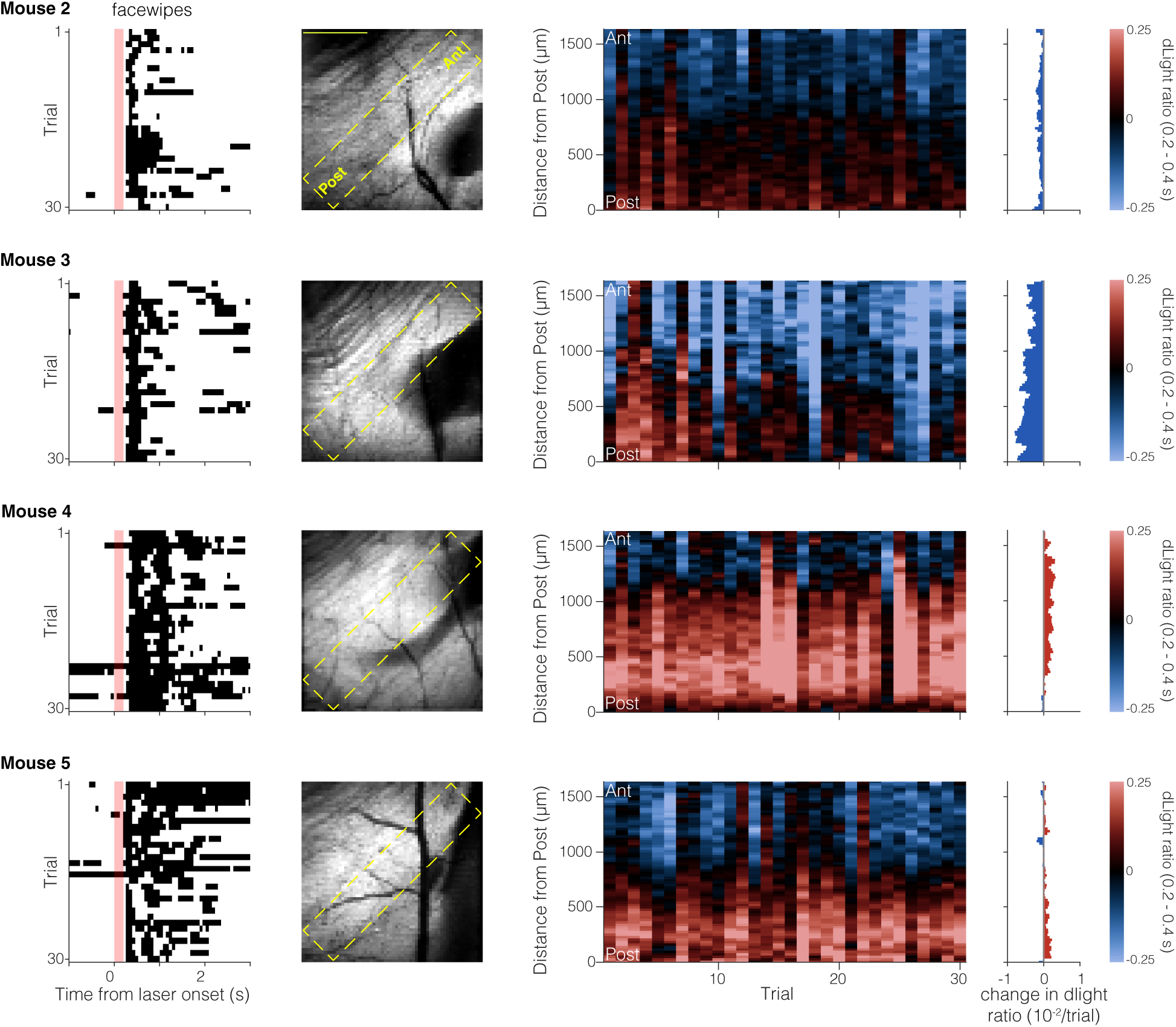
Stable responses to aversive heat stimulation across trials. *Left,* Raster plots aligned to heat onset showing face-wipe responses, defined by detected paw keypoint coordinates at or above a threshold set to 0.8 times the median lower-mouth y-coordinate across the recording. *Middle left,* Mean full-FOV images, indicating the diagonal strip used to generate kymographs. Data were acquired for 64 × 64 pixels at ≈205 Hz. Scale bar, 500 µm. *Middle,* Kymographs of mean dLight ratios from 0.2 – 0.4 s after heat onset across trials. *Middle right,* Trial-to-trial rate of change in dLight ratio along the posterior–anterior axis. For each distance bin, the mean 0.2–0.4 s dLight ratio was fit against trial number (1–30) by least-squares linear regression; horizontal bars show the resulting slopes for each anterior-posterior distance (red, increasing; blue, decreasing). Slopes are expressed as change in dLight ratio per trial (×10⁻²/trial). *Right,* color scale for the kymographs.

## Notes

### Competing Interest Statement

The authors have declared no competing interest.

